# Accumulation dynamics of defective genomes during experimental evolution of two betacoronaviruses

**DOI:** 10.1101/2024.02.06.579167

**Authors:** Julia Hillung, María J. Olmo-Uceda, Juan C. Muñoz-Sánchez, Santiago F. Elena

## Abstract

Virus-encoded replicases often generate aberrant RNA genomes, known as defective viral genomes (DVGs). When coinfected with a helper virus providing necessary proteins, DVGs can multiply and spread. While DVGs depend on the helper virus for propagation, they can disrupt infectious virus replication, impact immune responses, and affect viral persistence or evolution. Understanding the dynamics of DVGs alongside standard viral genomes during infection remains unclear. To address this, we conducted a long-term experimental evolution of two betacoronaviruses, the human coronavirus OC43 (HCoV-OC43) and the murine hepatitis virus (MHV), in cell culture at both high and low multiplicities of infection (MOI). We then performed RNA-seq at regular time intervals, reconstructed DVGs, and analyzed their accumulation dynamics. Our findings indicate that DVGs evolved to exhibit greater diversity and abundance, with deletions and insertions being the most common types. Notably, some high MOI deletions showed very limited temporary existence, while others became prevalent over time. We observed differences in DVG abundance between high and low MOI conditions in HCoV-OC43 samples. The size distribution of HCoV-OC43 genomes with deletions differed between high and low MOI passages. In low MOI lineages, short and long DVGs were most common, with an additional cluster in high MOI lineages which became more prevalent along evolutionary time. MHV also showed variations in DVG size distribution at different MOI conditions, though less pronounced compared to HCoV-OC43, suggesting a more random distribution of DVG sizes. We identified hotspot regions for deletions that evolved at high MOI, primarily within cistrons encoding structural and accessory proteins. In conclusion, our study illustrates the widespread formation of DVGs during betacoronavirus evolution, influenced by MOI and cell- and virus-specific factors.

## 1. Introduction

Owed to their high mutation rates, rapid replication and large population sizes, RNA viruses quickly adapt to novel hosts and trigger epidemics by crossing species barriers (Duffy, Shackelton and Holmes 2008; Sanjuán et al. 2010; Belshaw, Sanjuán and Pybus 2011). However, many mutations during infection lead to aberrant genomes that cannot complete the infectious cycle. *In vitro*, these incomplete genomes, known as defective viral genomes (DVGs), accumulate in virus populations when the virus is passaged repeatedly at high multiplicity of infection (MOI) (Stampfer, Baltimore and Huang 1971; Huang 1973; Felt et al. 2022). Some DVGs, called defective interfering particles (DIPs), can be encapsidated and transmitted when co-infected with a wild-type virus, interfering with its replication (Barret and Dimmock 1984; Roux, Simon and Holland 1991; Smith et al. 2016). This interference occurs on multiple levels, including resource competition, immune response activation, and sometimes, the survival of infected cells (Huang and Baltimore 1970; Huang 1973; Roux, Simon, and Holland 1991; Smith et al. 2016). Although specifics of DIP generation and regulation in RNA viruses are still not fully understood, their role in modulating viral infections is well-established (Rezelj, Levi, and Vignuzzi 2018; Vignuzzi and López 2019).

Deletions are the most common type of DVGs among RNA viruses (Di Gioacchino et al. 2022; Aguilar Rangel et al. 2023). They are thought to form through homologous recombination due to observed homology in specific regions and RNA structures (Jennings et al. 1983; Saira et al. 2013; Poirier et al. 2016). Copy-back (cb) and snap-back (sb) genomes are also prevalent DVG classes in negative-sense RNA viruses. These genomes have loops with complementary ends (Li and Pattnaik 1997; Sun et al. 2019).

Coronaviruses have the largest known RNA virus genomes, ∼30 kb in length (Brian and Baric 2005). They encode nonstructural proteins involved in viral RNA synthesis and interact with host cell functions. Replication generates a set of subgenomic mRNAs, all sharing common 5’ leader and a 3’ terminal sequences (Yang and Leibowitz 2015). Specific regions, such as the leader sequence and a portion of the replicase gene, contribute to efficient replication and transmission. Additionally, *cis*-acting elements like the transcriptional regulatory sequences (TRS) are crucial for transcription (Kim et al. 2020). In most described coronavirus’ DIPs, the combination of both internal and terminal genome sequences enables the formation of secondary and higher-order structures essential for their function as replication signals (Repass and Makino 1998). Deletion mapping of murine hepatitis virus (MHV) DIPs reveals the requirement of an internal and discontinuous sequence for replication (Lin and Lai 1993). The 3’ proximal coding regions, including the N cistron, can tolerate substantial alterations, suggesting they are not part of the 3’ *cis*-acting elements (Liu, Johnson and Leibowitz 2001; de Haan et al. 2002; Goebel, Taylor and Masters 2004; Liu et al. 2007). Efficient genome packaging into virions requires specific RNA signals, primarily located at the terminal sequences (Sethna, Hung and Brian 1989; Hofmann, Sethna and Brian 1990; van der Most, Bredenbeek and Spaan 1991; Fosmire, Hwang and Makino 1992; Zhao, Shaw and Cavanagh 1993).

To investigate the dynamics of DVG accumulation in betacoronaviruses infecting highly susceptible cells, we conducted an evolution experiment by performing undiluted passages of (*i*) the human coronavirus OC43 (HCoV-OC43) in baby hamster kidney cells (BHK-21) and human large intestine carcinoma cells (HCT-8), and (*ii*) of MHV in murine liver cells (CCL-9.1). We anticipated that high MOI would promote the accumulation of DVGs over passages due to frequent co-infection with WT viruses. As a control, we also carried out parallel evolution experiments at low MOI. We replicated both MOI conditions and virus-cell type combinations three times. After experimental evolution, we employed high-throughput RNA-seq and bioinformatic tools to reconstruct DVG populations in the ancestral virus and in samples at four equidistant evolutionary passages.

## 2. Results

### 2.1. Experimental evolution of betacoronavirus DVGs populations

Viral load was evaluated each passage and dilutions adjusted to ensure that, on median, MOIs remained within two wide but disjoint intervals that can broadly be defined as high and low MOIs, respectively (Supplementary Table S1). Notice that what is defined as high or low MOIs depends on the particular combination of virus and cell type, although in all cases, the log_2_-fold difference between high and low MOIs was consistently in the range 10 - 20.

The time-series data for viral load are shown in Fig. 1. Individual lineage series were fitted to ARIMA models and the best-fitting one determined according to the minimum BIC criterium (Supplementary Table S2). In each model, the slope parameter defines the rate of evolution per passage. The average rates of evolution for each combination of factors are also displayed in Supplementary Table S2. In the case of HCoV-OC43 in BHK-21 cells, the average rate was significantly positive at high MOI (runs test, *P* < 0.001) but not different from zero at low MOI (runs test, *P* = 1.000). On average, rates were 13% faster at high MOI. In the case of HC-OC43 in HCT-8 cell, both rates of evolution were significantly negative (runs tests, *P* < 0.001 in both cases). Rates were, on average 47% faster at high MOI, although the difference was not significant (Mann-Whitney, *P* = 0.719). Finally, MHV in CCL-9.1 cells showed no difference from zero at high MOI (runs test, *P* = 1.000) but significantly negative at low MOI (runs test, *P* < 0.001), declining 567% faster at low MOI (Mann-Whitney, *P* < 0.001).

**Figure 1.**
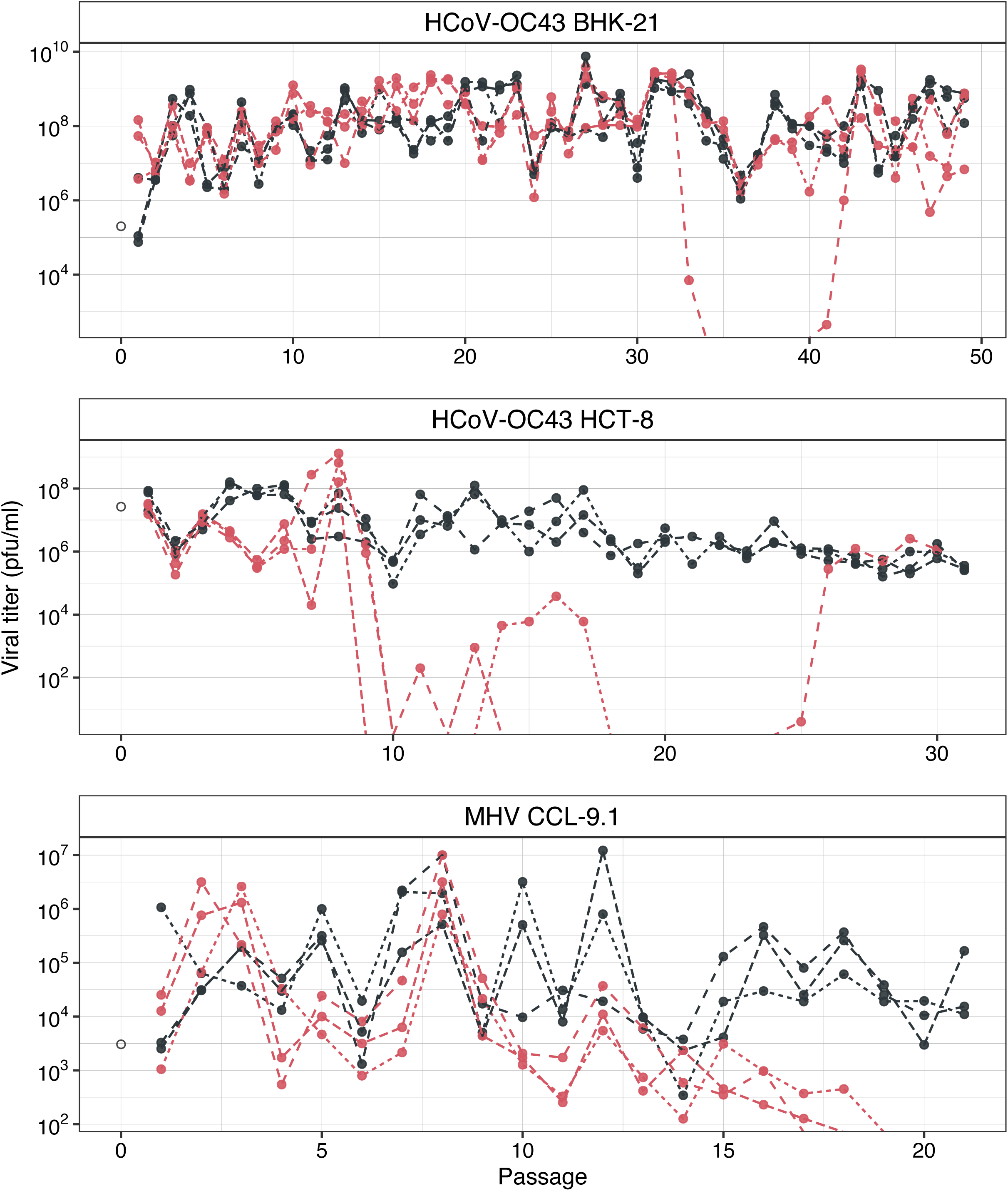
Evolution of viral loads at high and low MOIs. Upper panel: HCoV-OC43 in BHK-21. Middle panel: HCoV-OC43 in HCT-8. Lower panel MHV in CCL-9.1. Black dots and lines: high MOI; red: low MOI. Empty dot: ancestral virus (stock).

An overall decrease in viral load in most cases could be explained by two alternative hypotheses. Firstly, it could be attributed to the progressive accumulation of DVGs that interfere with the replication of wild-type viruses (Felt et al. 2022; Aguilar Rangel et al. 2023; Zhou et al. 2023). This effect would be stronger at high MOI, which seems not to be a general case in Fig. 1. Secondly, it could be a consequence of strong transmission bottlenecks that turns on Muller’s ratchet, leading to the fixation of deleterious mutations and, consequently, fitness declines, as well established for other RNA viruses (Clarke et al. 1993; Duarte et al. 1993; Novella et al. 1995; Elena et al. 1996; de la Iglesia and Elena 2007). To further explore the role of DVGs in these dynamics, we performed RNA-seq from the supernatants of infected cells from each lineage at four equidistant evolutionary time points, as well as from the corresponding ancestral viruses, in order to reconstruct the DVG populations at each sample.

### 2.2. Evolution of DVG richness and abundance

Since the sequencing was not strand-specific, in the following analyses we will not distinguish between DVGs found in the negative or positive sense RNA strains. In addition, prior to all the evolutionary relevant analyses, canonical subgenomic RNAs (sgRNAs) were filtered out by excluding any reconstructed DVGs whose junctions agreed with the expected coordinates of sgRNAs within a range of ±10 nucleotides. The percentages of sgRNAs observed in each evolving lineage are shown in Supplementary Fig. S1. Interestingly, these percentages were affected by MOI, virus species and host cell type. In the case of HCoV-OC43, 51.35% less sgRNAs relative to total DVGs were observed at high MOI than at low MOI in BHK-21. This reduction was 41.44% in HCT-8. These observations suggest that, for a given amount of sgRNAs, more DVGs were accumulating at high MOIs. In contrast, the trend was the opposite for MHV in CCL-9.1: 19.26% more sgRNAs per total DVGs at high than at low MOIs.

Next, we examined the dynamics of DVG generation and accumulation during the evolution experiments. Fig. 2 shows two different measures of DVG diversity along time. Firstly, the number of unique DVGs per 10^5^ viral reads (RPHT) (Fig. 2A), which represents diversity in terms of different DVG types observed. Secondly, the total RPHT per DVGs (Fig. 2B), a measure of the total amount of DVGs relative to absolute viral accumulation. In all lineages, a common trend was observed: an increase in DVG richness from the initial to intermediate stages of viral evolution (Fig. 2A), with no significant effect of MOI (Wilcoxon paired-samples tests, *P* ≥ 0.382 in all three cases). Throughout the course of the evolution experiment, some disparities were noted between parallel replicates, but the overall pattern for HCoV-OC43 in both cell types was a reduction in richness after the initial surge, maintaining a relatively stable level until the end of the experiment, with no discernible differences observed between both cell types. For MHV at high MOI, DVG richness increased up to passage nine and then remained stable until the end of the experiment. However, at low MOI, MHV displayed differences between parallel lineages, with the number of DVG reads either increasing or decreasing after passage nine, depending on the lineage. The shifts in richness at different passages were more remarkable for HCoV-OC43 in BHK-21 and for MHV in CCL-9.1 cells.

**Figure 2.**
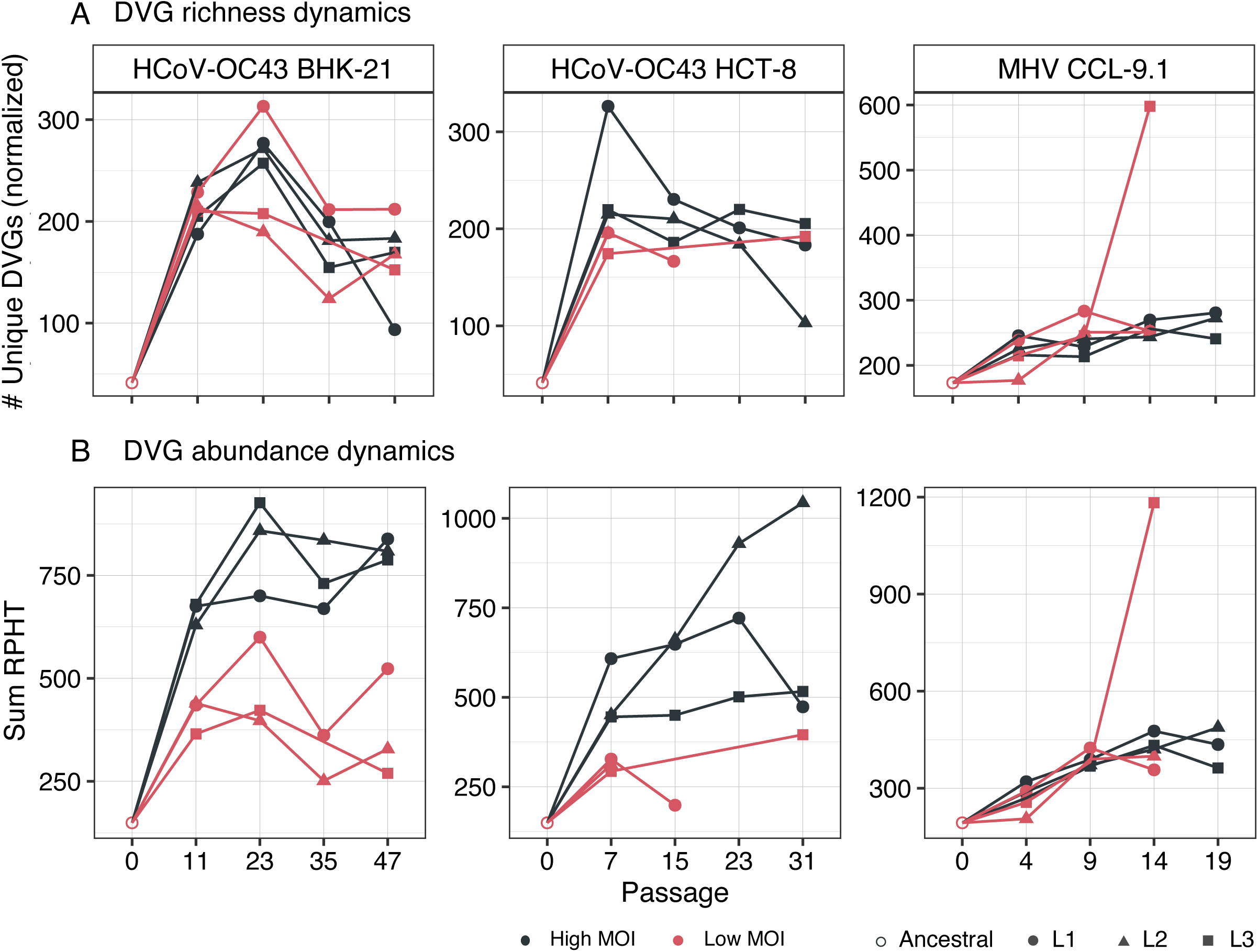
Evolution of richness and abundance of DVGs. **(A)** DVGs richness, as the number of unique DVGs relative to the total viral RPHT. **(B)** DVGs abundance, estimated as the total number of DVG read counts relative to the total viral reads in the sample, also RPHT. Black colour represents high MOI while red represents low MOI. Different lineages are presented by different symbols. When a lineage was extinct, no point is shown.

Furthermore, the abundance of DVG counts increased to varying degrees at the onset of evolution in all lineages (Fig. 2B). After the initial passages in low MOI lineages, the abundance of detected DVGs tended to decline, while in high MOI lineages, DVG abundance remained relatively steady or even slightly increased. Interestingly, notable disparities were noted in the accumulation of DVGs between high and low MOIs for HCoV-OC43 in both cell types (Wilcoxon paired-samples tests, *P* = 0.022 and *P* = 0.041, respectively), with DVGs consistently being more abundant in lineages evolved at high MOI. In contrast, no significant differences in terms of the total number of accumulated DVGs were observed for MHV lineages evolved at high and low MOIs (Wilcoxon paired-samples test, *P* = 1.000).

Next, we tested whether DVG richness and abundance were correlated, that is, whether more abundance would contain more different DVG types. The alternative being abundance resulting from the accumulation of few different DVGs. Spearman correlations were significant for HCoV-OC43 infecting BHK-21 at low MOI (*r_S_* = 0.882, 13 d.f., *P* = 0.001) as well as for MHV at both MOIs (low: *r_S_* = 0.900, 10 d.f., *P* = 0.002; high: *r_S_* = 0.776, 13 d.f., *P* = 0.005) but not in the other three cases. These observations suggest that viral populations containing more DVGs may also contain more diverse types, but not in all instances.

### 2.3. Reconstructed DVG types

DVG classes considered in the reconstruction include deletions, insertions, and cb or sb genomes (hereafter all referred to as cb) at 5’ or 3’ ends. Richness was evaluated as in Section 2.2 but now distinguishing between the four types of DVGs; estimates are shown in Fig. 3A. Firstly, we asked whether the two viruses show differences in the number of unique DVGs types. A Scheirer-Ray-Hare (SRH) two-ways non-parametric ANOVA test showed highly significant differences between the two viruses (*P* < 0.001), with HCoV-OC43 generating, on average, 47% more different DVGs than MHV. Likewise, the overall distribution of different DVGs also differed among the four classes of DVGs, with deletions being the most abundant type, followed by insertions and with 3’ and 5’ cb being the less common (*P* < 0.001). Interestingly, these differences among the four classes were consistent for the two viruses (*P* = 0.442). Secondly, we interrogated the HCoV-OC43 data for differences among cell types. In this case, the SRH test found no differences in the number of different DVG types among the two cells (*P* = 0.093), although the distribution of the four DVG types was strongly biased by deletions and insertions (*P* < 0.001). Thirdly, we tested whether MOI had an overall effect on the number of different DVGs generated. In this case, the SRH test also found significant differences (*P* < 0.001) between lineages evolved at high and low MOIs, with the former containing 74% more variants than the latter. The rank-order of the four classes of DVGs was not affected by MOI (*P* = 0.894). Abundance was evaluated as in Section 2.2 but now distinguishing between DVG types; estimates are shown in Fig. 3B. Following the same logic than in the previous paragraph, we firstly sought for an effect of virus species on DVG abundance. In this case, the SRH test found no significant differences among the two viruses in the abundance of DVGs (*P* = 0.575). Overall, deletions were the most variable class, followed by insertions and 5’ and 3’ cb (these two at approximately the same variability) (*P* < 0.001); these differences were not affected by the viral species (*P* = 0.093). Secondly, we found no significant differences among the two cell types in the abundance of DVGs generated from HCoV-OC43 genome (*P* = 0.835), nor an effect on the distribution of the four classes of DVGs (*P* = 0.994). Finally, differences in MOI had an overall effect on DVGs abundance, being 35% more abundant in lineages evolved at high MOI (*P* < 0.001), although this effect did not translate into a change in the proportion of the four classes of DVGs (*P* = 0.856).

**Figure 3.**
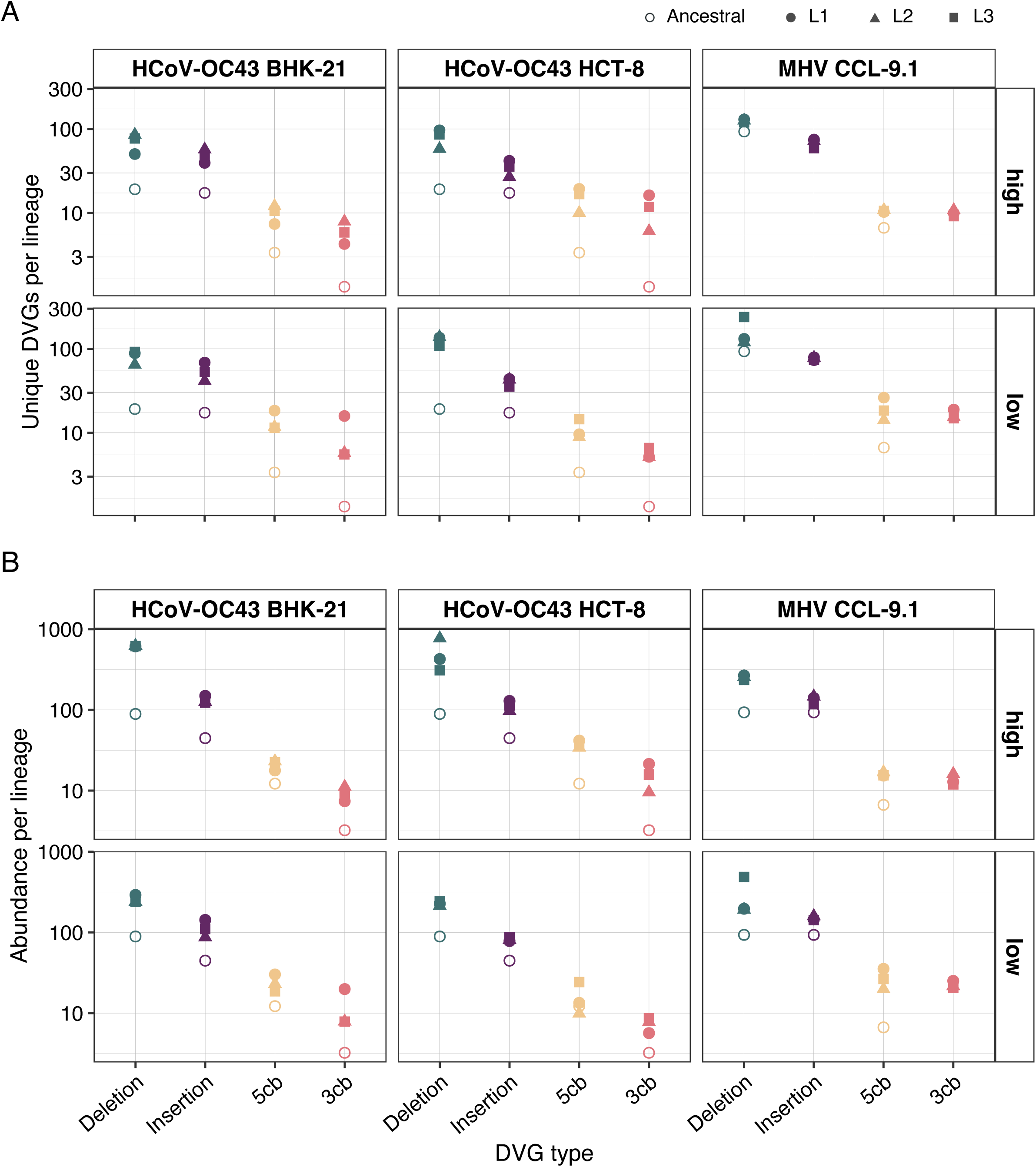
Diversity of DVG classes detected in each lineage of experimental evolution. **(A)** Richness represented as the normalized number of unique DVGs present in each lineage. **(B)** Abundance represented as the normalized number of reads supporting the DVG founded in each lineage. Empty points represent the stock sample (passage 0), the values for the stock in HCoV-OC43 are the same for BHK-21 and HCT-8 at both MOIs. The MHV ancestral is the same for both MOIs in CCL-9.1.

### 2.4. The distribution of deletions sizes

Next, we honed in on deletions for our analysis, as they were the most common type of DVGs. We examined the sizes of defective genomes with deletions at each sampled time point. One-base deletions were predominant in all the lineages. Thus, to uncover the pattern of remaining deletion sizes, we filtered out DVGs with deletions shorter than 50 nucleotides.

The evolution of the distribution of deletion sizes, accounting for their corresponding abundance, is shown in Fig. 4A. The size distribution of HCoV-OC43 genomes with deletions differed between high and low MOI passages in both cell types, with bigger deletions being more common at high MOI (Fig. 3A, black bars). Notably, certain deletion sizes at high MOI became predominant throughout the evolution (BHK-21: ∼12.4 kb, ∼22.5 -30.7 kb; HCT-8: ∼13.6 kb). HCoV-OC43 evolved to cell-specific size distributions, although some lengths were common to both cell types. At low MOI (red bars in Fig. 4A; for a more detailed presentation, see Supplementary Fig. S2), large deletions sizes ∼30 - 31 kb are predominant for HCoV-OC43, followed by ∼18 kb in BHK-21 and ∼18 - 20 kb in HCT-8 cells. The 30,698-nucleotides long DVG is widely present across all the lineages and MOIs of HCoV-OC43 infected cells but it was also present in the viral stock.

**Figure 4.**
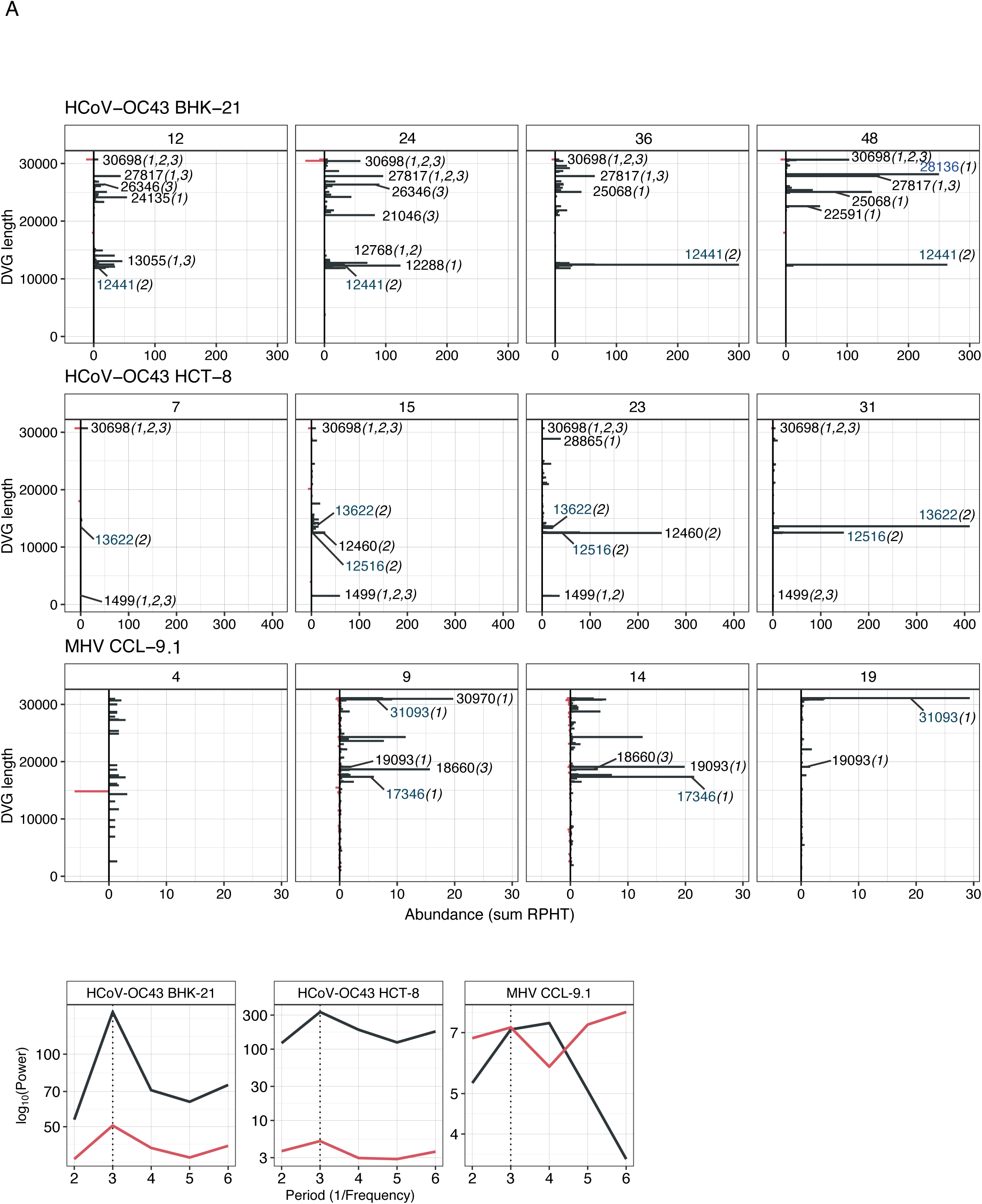
Distribution of deletion sizes. **(A)** Changes in the size distribution of deletions along the evolution experiments. Size distribution of genomes with deletions across different evolutionary time points (upper box of each panel, passage numbers), taking into account the variability and abundances of the reconstructed deletions. Replicates of lineages were combined into a single diagram. Numbers indicate some predominant sizes, between parenthesis the lineages in which they were found. Blue numbers highlight the lengths that gain predominance with the evolution. Genomes with deletions of fewer than 50 nucleotides were excluded from the visualization. **(B)** Lomb-Scargle periodogram of deletion sizes. All deletions > 3 nucleotides where include in the frequency analysis.

For MHV there were also differences in the distribution of deletion sizes between high and low MOIs, although less obvious than shown for HCoV-OC43. The size of the most frequent deletions at high MOI were ≥ 17 kb (Fig. 3A, black bars), with varying maxima to different time points along viral evolution. The last passage showed a clear predominance of the longest deletions (∼31.1 kb, 19.1 kb). However, the distribution of deletion sizes at low MOI lacked of clear abundance pattern (Supplementary Fig. S2).

From these analyses, we can conclude that although the generation of short deletions is more likely, the occurrence of very long deletions is also common. The size distribution of deletions is highly dynamic and underwent changes throughout the course of the evolution experiment, with certain lengths being pervasively favored. This dynamism is influenced by the combination of viral species and cell type, being the viral species the most influential factor.

Aguilar Rangel et al. (2023) proposed the idea that the triplet periodicity of the genetic code must place a constraint on deletions in coding sequences. To test this hypothesis, we evaluated whether a characteristic multiplicity existed in the distribution of deletions size. A Lomb-Scargle periodogram (an adaptation of the classic Fourier transformation for unevenly sampled data; VanderPlas 2018) of deletion sizes revealed a robust tri-nucleotide periodicity in the cistrons (Fig. 4B). HCoV-OC43 data in both cell sizes and MOIs show a characteristic frequency multiple of three (a strong signal at period three and a weaker one at period six). No obvious periodicity has been observed for MHV at high MOI but a weak signal at period three and a stronger one at period six appears to exist (Fig. 4B). This analysis suggests that deletions more likely to disrupt the translation reading frame did not accumulate as much as those that keep it, suggesting that DVG size was evolving under purifying selection.

### 2.5. Factors influencing the occurrence of deletions

Next, we conducted an analysis to determine if specific domains of the viral genome were more prone to act as break points (BP or start) and rejoining points (RP or end) for deletions owed to their sequence or involvement in structural elements. Firstly, we hypothesized that the emergence of deletions could be linked to the very same mechanisms involved in sgRNA formation. These mechanisms in coronavirus genome involve TRS. To test this possibility, we calculated the frequency of each genomic position as BP or RP of a deletion (Fig. 5A). Deletions shorter than three nucleotides, were excluded hereafter based on the consideration that very small deletions, such as sequencing errors, must follow a different distribution compared to larger, biologically relevant events. At this stage, we want to restress out that canonical sgRNAs were also filtered out before the analysis to avoid spurious results.

**Figure 5.**
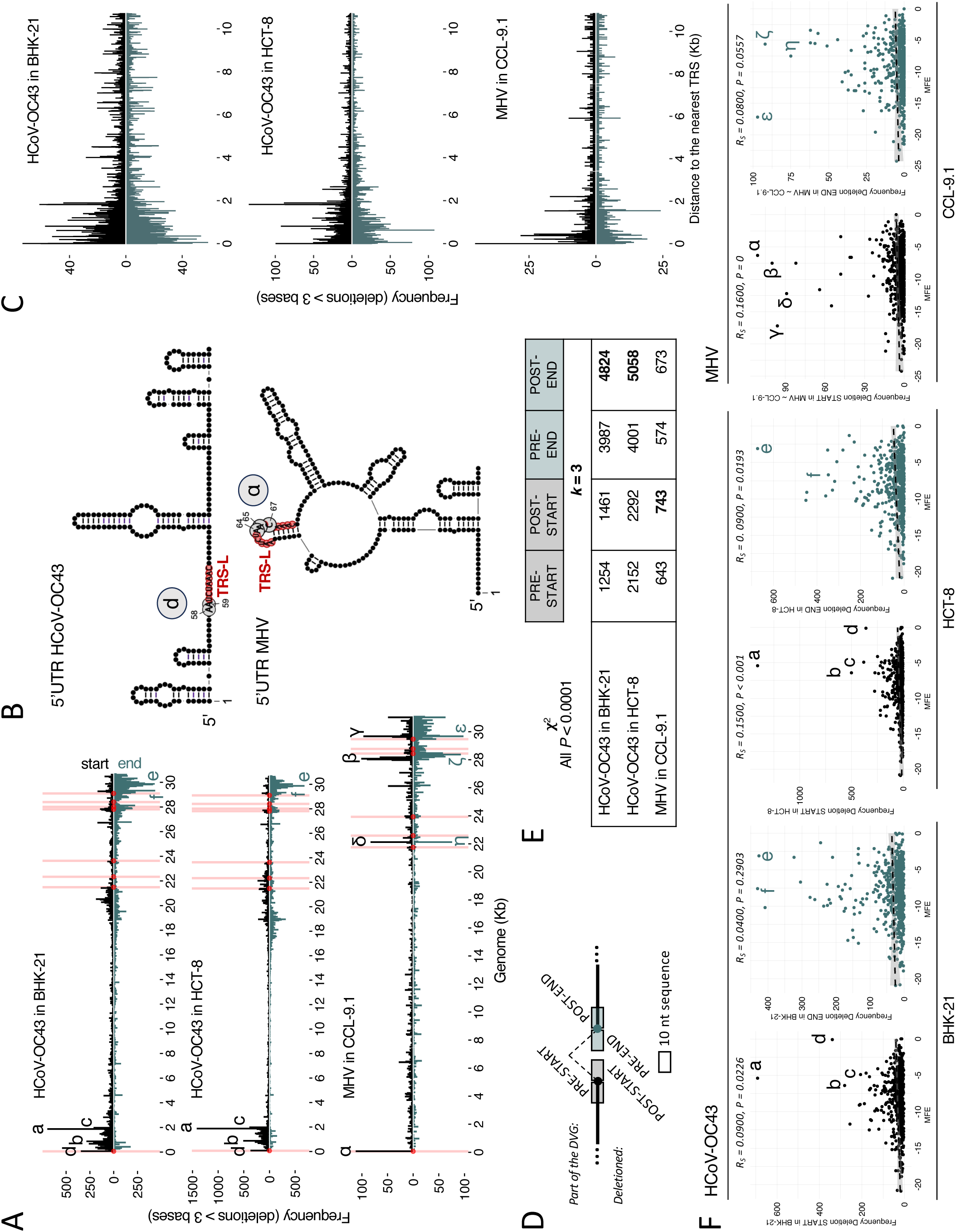
Genomic determinants of DVG formation. **(A)** Frequencies of BP and RP of deletions > 3 at each genome positions (in kb). Each bar of the histogram represents a 50-bases window. The upper part of the diagrams indicates start points, while the lower part does so for end points. Specific TRS positions are highlighted with pink bars and with circles at the base of the start and end points. The most frequently affected areas are indicated with letters a-f for HCoV-OC43 and α-η for MHV. HCoV-OC43 ranges: a = 1850 - 1900; b = 850 - 900; c = 1950 - 2000; d = 50 - 100; e = 29,850 - 29,900; and f = 29,800 - 29,850. MHV ranges: α = 50 - 100; β = 28,050 - 28,100; γ = 29,650 - 29,670; δ = 28,000 - 28,050, ε = 29,650 - 29,700, ζ = 28,350 - 28,400, and η = 2100 - 22,150. **(B)** 5’UTR structure prediction and localization of one of the most frequent start points (d in HCoV-OC43 and α in MHV) of deletions; TRS-L highlighted in red. **(C)** The distance in nucleotides from each deletion’s BP (above) and RP (below) to the nearest TRS (in kb). **(D)** Schematic representation of DVG junction with pre- and post-BP and RP sequences as studied in the *k*-mer analysis. **(E)** Results from the χ^2^ test of 3-mer distribution between the sequences around the deletion compared to those found in the complete genome. Sequences of 10 nucleotides before the BP (pre-BP), 10 after the RP (post-RP), 10 after the BP (post-BP), and 10 before the RP (post-RP) were analyzed. All the results are significant (*P* < 0.001), with the highest χ^2^ value of each case bolded. **(F)** Correlation between frequency of BP (black) or RP (green) points with the minimum free energy (MFE) predicted for the 50-bases window secondary structure.

In HCoV-OC43, BP accumulated within the first 2 kb of the genome, while the more frequent RP accumulated within the last 2 kb (Fig. 5A), with additional likely sites around 415 nucleotides from the 3’ end in BHK-21 and 765 and 1215 in HCT-8. Interestingly, the most likely BP d is two nucleotides upstream from the TRS-L and few nucleotides before a large structural hairpin in 5’UTR (Fig. 5B). This observation was consistent for both cell types. In the case of MHV, the most likely BP was labeled as α, and it lied in the middle of TRS-L and the loop of a hairpin structure (Fig. 5B). The density of potential BP and RP in MHV was also high at the last third of the genome for BP and RP, but highly frequent sites were also found in the TRS-L sequence (67 nucleotides), at 970 and at 5975 nucleotides for BP and at 996 and 5996 nucleotides for RP (Fig. 5A).

To further explore into the proximity of BP and RP to canonical the breakpoints for sgRNA generation (*i.e*., TRS), we calculated the distance in nucleotides from each deletion’s BP (above) and RP (below) to it nearest TRS (Fig. 5C). The frequency distribution at various distances from TRS (∼0 - 11 kb) appeared relatively symmetric for both BP and RP (Fig. 5C). The most frequent recombination points in all cell types, but particularly in BHK-21, were concentrated within a range of 2 kb around TRS (where the a - c hotspots were also concentrated; Fig. 5A), showing a consistent decrease in frequencies with increasing distance from TRS (Fig. 5C).

Next, we explored whether the likelihood of being BP and RP can be attributed to specific sequences. The richness in A/U sequences around breakpoints was documented previously for influenza virus (Elshina and te Velthuis, 2021) and previous studies with coronaviruses pointed the UUG triplet as the preferred sequence for spanning junction start positions and a significant A increase in the end positions (Gribble et al., 2021). Following that, we analyzed the distribution of nucleotides (*k*-mer analysis) of the surroundings of the recombination points (Fig. 5D). Fig. 5E shows the results of the χ^2^ test supporting distinct 3-mer distributions between sequences around the deletion coordinates and those found in the entire genome (*P* < 0.001 in all cases). Analysis of shorter *k*-mers produced identical results (not shown). In addition to all nucleotide’s distributions being significantly different at the recombination coordinates, we observed in HCoV-OC43 that, irrespective of the cell type, the most notable changes in the *k*-mer distribution were observed in the sequences surrounding the RP. In MHV, the highest χ^2^ values were in the sequences downstream the BP (Fig. 5D).

Inspired by the results shown in Fig. 5B associating highly frequent BP with RNA secondary structures, we tested whether the likelihood of being BP or RP could be associated with the complexity of secondary structures. We assumed, that the lower the minimum free energy (MFE), the more complex the RNA structure (Zuker 1989) and the more likely the RNA polymerase will jump out the template and rejoin somewhere else. As shown in Fig. 5F, significant Spearman correlations were observed between the frequency of BP and MFE for both viruses and, in the case of HCoV-OC43, both cell types (in all cases, *P* :: 0.023). Regarding RP, a significant correlation was only found for HCoV-OC43 in HCT-8 (*P* = 0.019). Qualitatively, regions with the highest deletion frequency tended to have intermediate MFE values. This suggests that deletions occurred more frequently in structured regions, but not necessarily in highly complex structures.

### 2.6. Cistrons affected by deletions

Now our focus moved to explore whether deletions evenly affected all cistrons or were preferentially found in certain cistron. To achieve this, we calculated the density of deletions per cistron by dividing the number of deletions found in a given cistron by its length. Data are shown in Fig. 6. These data were then fitted to a generalized linear mixed model (GLMM) in which cell type, MOI and cistron were treated as fixed effects and lineage (nested factor within the interaction of the fixed factors) and passage (within-individual repeated measures) as random ones, and a Gaussian distribution and identity link function. Focusing in HCoV-OC43, no differences among cell types (main or in combination with other fixed factors) were found (χ^2^ :: 1.971, *P* := 0.923). MOI had a net effect, with density across cistrons being 178% higher at high MOI (χ^2^ = 55.961, 1 d.f., *P* < 0.001). Also, significant overall differences existed among passages (χ^2^ = 36.509, 6 d.f., *P* < 0.001), with the density of deletions across cistrons increasing with evolutionary time. Highly significant differences were found among cistrons (χ^2^ = 126.321, 7 d.f., *P* < 0.001), with ORF1ab and S showing the lowest density of deletions whilst E and ns12.9 shown the highest. Most interestingly, a significant interaction between MOI and cistron was observed (χ^2^ = 36.298, 7 d.f., *P* < 0.001), with a reduction in the magnitude of the differences between cistrons at low MOI.

**Figure 6.**
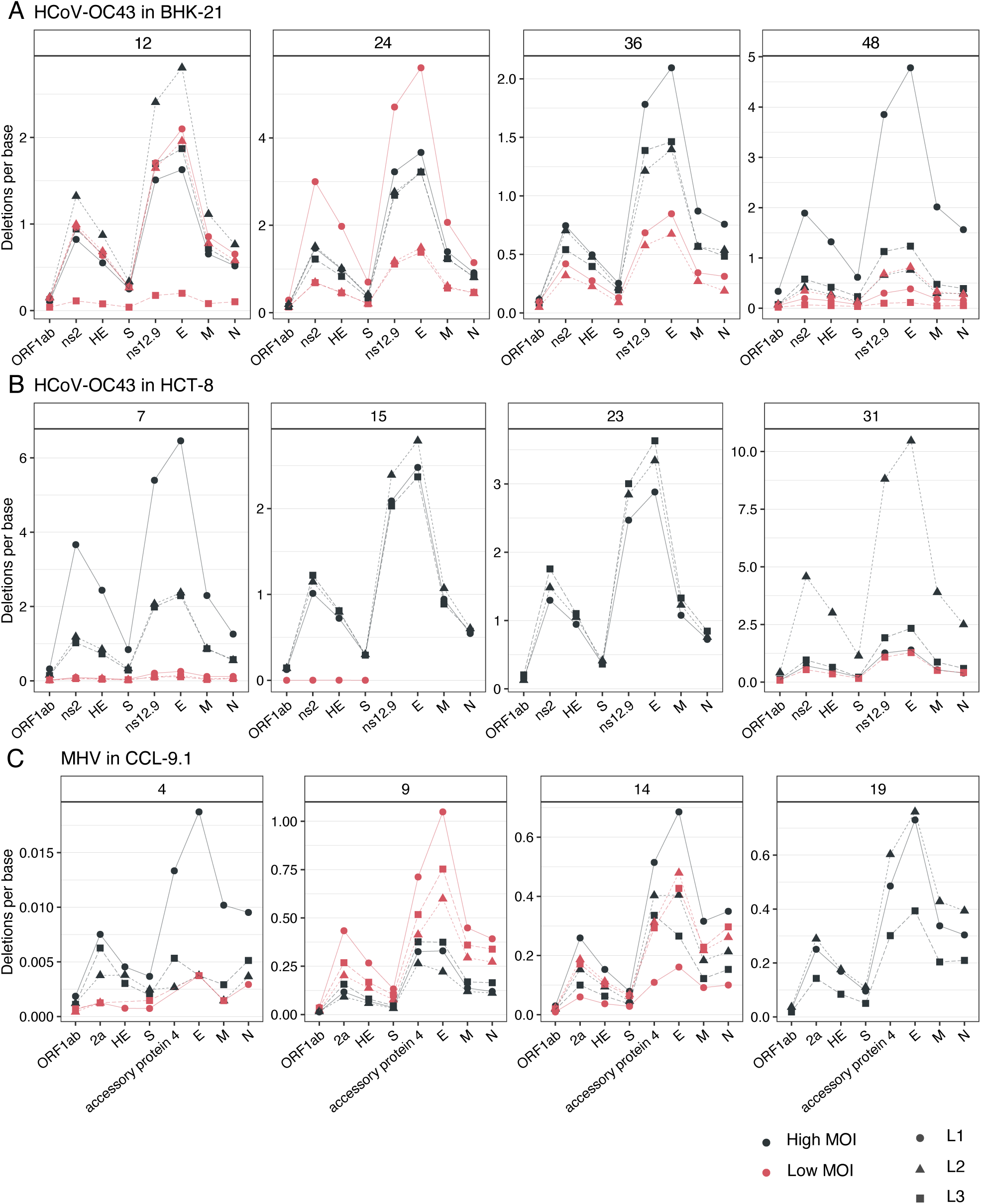
Observed frequency of deletions per cistron per base. Each panel corresponds to the indicated passage in the evolution experiment.

The results for MHV were somehow different. A significant increase in the overall deletion density along evolutionary time (χ^2^ = 88.730, 3 d.f., *P* < 0.001) was observed. Significant differences among cistrons existed (χ^2^ = 75.268, 5 d.f., *P* < 0.001), again with ORF1ab and S showing the lower deletion density and the auxiliary protein 4 and E showing the highest. All other factors and combinations were not significant.

### 2.7. A subset of DVGs persists along the evolution experiment

Next, we examined the temporal changes in the proportions of persistent DVGs during the experimental evolution. DVGs qualified as *de novo* if observed only in a single passage, while they were qualified as persistent if, after the first detection, were observed at subsequent passages of the same lineage. For each virus species and cell type independently, data in Fig. 7A were fitted to a GLMM with MOI and type of DVG treated as fixed effects and lineage (nested factor within the interaction of the fixed factors) and passage (within-individual repeated measures) as random ones, and a Gamma distribution and log link function. In the case of HCoV-OC43 in BHK-21, as the number of passages increases, the percentage of persistent DVGs rose consistently in BHK-21 cells for all DVGs types (χ^2^ = 18.830, 3 d.f., *P* < 0.001) regardless of the MOI (χ^2^ = 4.872, 3 d.f., *P* = 0.181). Interestingly, highly significant overall difference was found among DVGs types (χ^2^ = 515.509, 3 d.f., *P* < 0.001), with deletions and insertions being the most common type of persistent DVGs (consistent with the results shown in Section 2.3). Yet, the rate at which DVGs accumulated was the same for all types (χ^2^ = 8.859, 9 d.f., *P* = 0.450). Another interesting result of this analysis was the significant interaction between DVG type and MOI (χ^2^ = 13.705, 3 d.f., *P* = 0.003), due to a disproportionate larger reduction (57.68%) of deletions at low MOI (5.31% for 3’ cb, 4.39% for 5’ cb and 22.33% for insertions). However, the pattern observed for HCoV-OC43 in HCT-8 cells differed from that in BHK-21 cells. There was a net overall effect of MOI (χ^2^ = 14.602, 1 d.f., *P* < 0.001), with 65.16% more DVGs accumulating at high MOI, being this effect also dependent on the evolutionary time (χ^2^ = 6.332, 2 d.f., *P* = 0.042), with an increasing trend for all types of DVGs at high MOI but mostly declines at low MOI.

**Figure 7.**
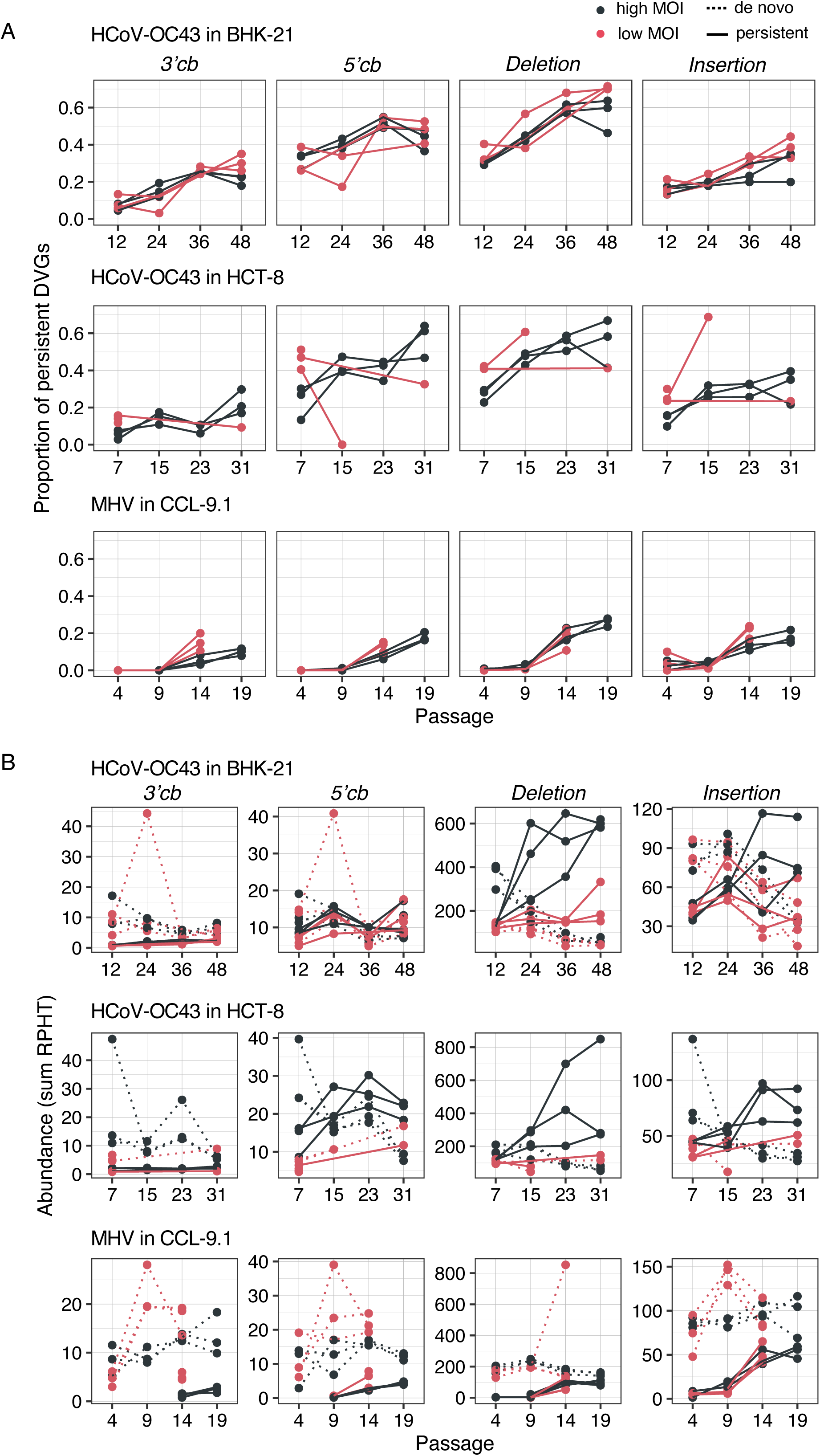
Transmission of persistent DVGs. **(A)** Proportion of persistent DVGs at different time points of experimental evolution, separated by DVG type. The proportion was calculated as the ratio of DVGs classified as persistent among all DVGs in the sample. **(B)** Abundance of persistent (solid lines) and *de novo* generated (dotted lines) DVGs at high (black) and low (red) MOIs, separated by DVG type. Evolutionary lineages are represented by shapes. RPHT: DVG reads per hundred thousand viral reads.

The case of MHV shown similarities and differences with HCoV-OC43 (Fig. 7A) as all the terms in the GLMM were highly significant. Among similarities, the proportion of persistent DVGs increased with the number of passages (χ^2^ = 72.091, 3 d.f., *P* < 0.001), although this increase depended both in the type of DVG and on MOI (χ^2^ = 85.477, 6 d.f., *P* < 0.001). Also, overall, the abundance of DVGs depended on their type, with deletions being the most common followed by insertion, 5’ and 3’ cbs (χ^2^ = 253.615, 3 d.f., *P* < 0.001). The most remarkable difference was that the overall effect of MOI went in the opposite direction: 24.15% more accumulation at low MOI. The increase in DVG abundance at low MOI might seem counterintuitive, but various factors, such as the possibility of secondary infections at low MOI (especially for MHV with a short infection cycle), could contribute to co-infection and the eventual transmission of DVGs.

We considered that the most parsimonious explanation for the temporal persistence of DVGs was that they were transmitted across passages. This assumption was supported by the occurrence of deletions being not random but associated with specific regions of the viral genome, suggestive of some sort of selection at play. However, we could not rule out the possibility that some deletions might pervasively emerge *de novo* at the same coordinates. To test this possibility, we calculated the relative abundance of reads categorized as persistent or as *de novo* at the various time points, distinguishing between different DVG types and MOI condition (Fig. 7B). This distinction was added as an additional fixed factor to the GLMM above and its effect on the abundance of the different types of DVGs evaluated. In the case of HCoV-OC43 in BHK-21 cells, this factor was significant by itself (χ^2^ = 26.159, 1 d.f., *P* < 0.001) as well as in interaction with all the other factors (in all cases, χ^2^ := 6.613, *P* :: 0.010). Three particularly relevant cases were: (*i*) the interaction of this factor with passage number (χ^2^ = 26.159, 1 d.f., *P* < 0.001): while de abundance of the *de novo* DVGs decreased, on average, 54.17% along the evolution experiment, transmitted DVGs had increased 114.36%. (*ii*) The interaction with MOI (χ^2^ = 6.613, 1 d.f., *P* = 0.010). At high MOI, transmitted DVGs were 46.01% more abundant than at low MOI, while *de novo* ones were only 19.87% more, suggesting than high MOI favors the transmission of already existing DVGs. And (*iii*) the interaction with DVG type (χ^2^ = 250.006, 3 d.f., *P* < 0.001). While deletion DVGs were 146.50% more abundant among the transmitted class than among *de novo* class, all other types were less abundant in the transmitted (79.74% for 3’ cb, 5.99% for 5’ cb and 4.86% for insertions). When analyzing the patterns in HCT-8 cells, results were not fully congruent. For example, the interaction with MOI was not significant (χ^2^ = 0.403, 1 d.f., *P* = 0.525). The situation with MHV was similar to HCoV-OC43 in HCT-8 and we are not discussing it on detail.

### 2.8. Hotspots of BP and RP for persistent DVGs

In Sections 2.5 and 2.6, we demonstrated that recombination events leading to deleted genomes were not randomly distributed across the genome. Here, we explored whether specific hotspots were overrepresented among persistent DVGs, potentially linked to their viability in *trans*-complementation with the helper virus. Fig. 8 illustrates the coordinates of BP and RP for persistent DVGs across all evolutionary lineages. As reference, Fig. 8A and Fig. 8C show schematic representations of the viral genomes. Overall, the most frequent coordinates in HCoV-OC43 infecting BHK-21 (Fig. 8B top) aligned with the findings shown in Fig. 5A. Start points were notably concentrated within the first 2 kb at the 5’ end of the genome, with another densely populated region between positions 20 - 22 kb. Notably, BP exhibit minimal variability, while RP were more diverse, spanning positions 17 - 30 kb. This was particularly evident for the two vertical lines around positions 50 and 2000, corresponding to the previously characterized a and d hotspots (Fig. 5A). At low MOI, we observed fewer hotspots, with a high density within the first 2 kb, similar to high MOI (Fig. 8B top). However, the high-density points in the last third of the genome at high MOI were absent in the low MOI samples. In lineages evolved in HCT-8 (Fig. 8B bottom), persistent deletions condensed at the same hotspots as in BHK-21. A notable difference was observed: in HCT-8, the variability of BP increased at specific positions, forming a range of BP spanning positions 0 to 5 kb and another range around positions 20 to 24 kb (Fig. 8B bottom). These hotspots seemed to be specific to HCoV-OC43 in HCT-8 cells. At low MOI, the limited number of persistent DVGs hampered pattern recognition, but the identified spots aligned well with those sawn at high MOI.

**Figure 8.**
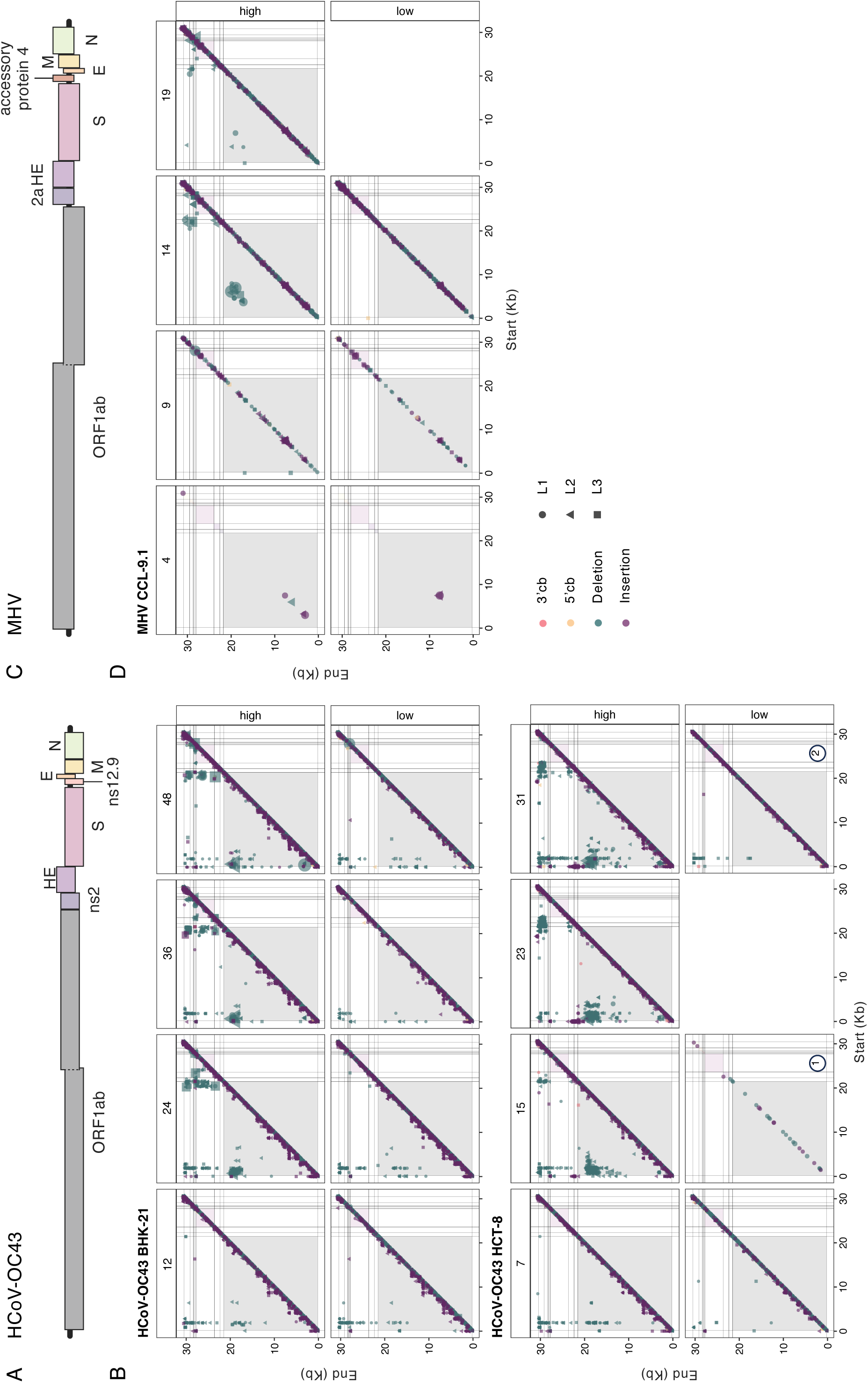
Distribution of persistent DVGs BP and RP. **(A)** Schematic representation of HCoV-OC43 genome. (**B)** Coordinates found for HCoV-OC43, at each passaged characterized, in both cell types. **(C)** Schematic representation of MHV genome. **(D)** As in panel B but for MHV. Coordinates from parallel lineages were joined into the same plot and represented by different symbols. Background colors indicate the cistrons as in panels A and C.

The number of spots for persistent DVGs was small for MHV at both high and low MOIs (Fig. 8D). However, two regions with accumulated coordinates for DVGs were observed at latter evolutionary passages. These hotspots spanned from positions 3 to 9 kb and from 16 to 22 kb.

### 2.9. Comparison of deletion composition among replicate lineages

Next, we aimed to assess the level of reproducibility in the generation of DVGs amongst lineages of the same virus evolved in the same cell host type. Fig. 9 depicts the counts of unique deletions (> 3 nucleotides) shared among the three independent lineages. A question that immediately arose was whether the count of shared unique DVGs among lineages differed from what would be expected based on the composition of each individual lineage. To illustrate this assessment, lets focus on HCoV-OC43 in BHK-21 at high MOI (Fig. 9A). The total count in lineage 1 was 4920 + 116 + 140 + 126 = 5302, 3831 for lineage 2, and 3465 for lineage 3. The count of unique DVGs observed exclusively in lineage 1 was 4920. The expected counts for this group can be calculated as (4920/5302)·(1 – 3532/3831)·(1 – 3142/3465)·(5302 + 3831 + 3465) ≈ 85. Using the same rationale, the expected counts could be computed for the other six groups displayed in the panel (BHK-21 low: ≈ 53; HCT-8 high: ≈ 248; HCT-8 low: ≈ 13; CCL-9.1 high: ≈ 14; CCL-9.1 low: ≈ 2). Applying a goodness-of-fit test, we discovered that observed counts significantly deviated from the expected values (χ² = 586,779.739, 6 d.f., *P* < 0.001). This deviation was primarily due to the deficit in counts in the three pairwise comparisons and, specially, in counts in the group shared by all three lineages. This finding suggested that most of the observed deletion DVGs were produced in a lineage-specific manner, while very few were produced in an almost deterministic manner and shared by all lineages. Similar qualitative outcomes were obtained for HCoV-OC43 in other scenarios (Fig. 9A and Fig. 9B), with a very large excess of linage-specific unique DVGs (χ² ≥ 15,940.544, 6 d.f., *P* < 0.001). Same results were found for MHV at both MOIs (χ² ≥ 71,675.329, 6 d.f., *P* < 0.001).

**Figure 9.**
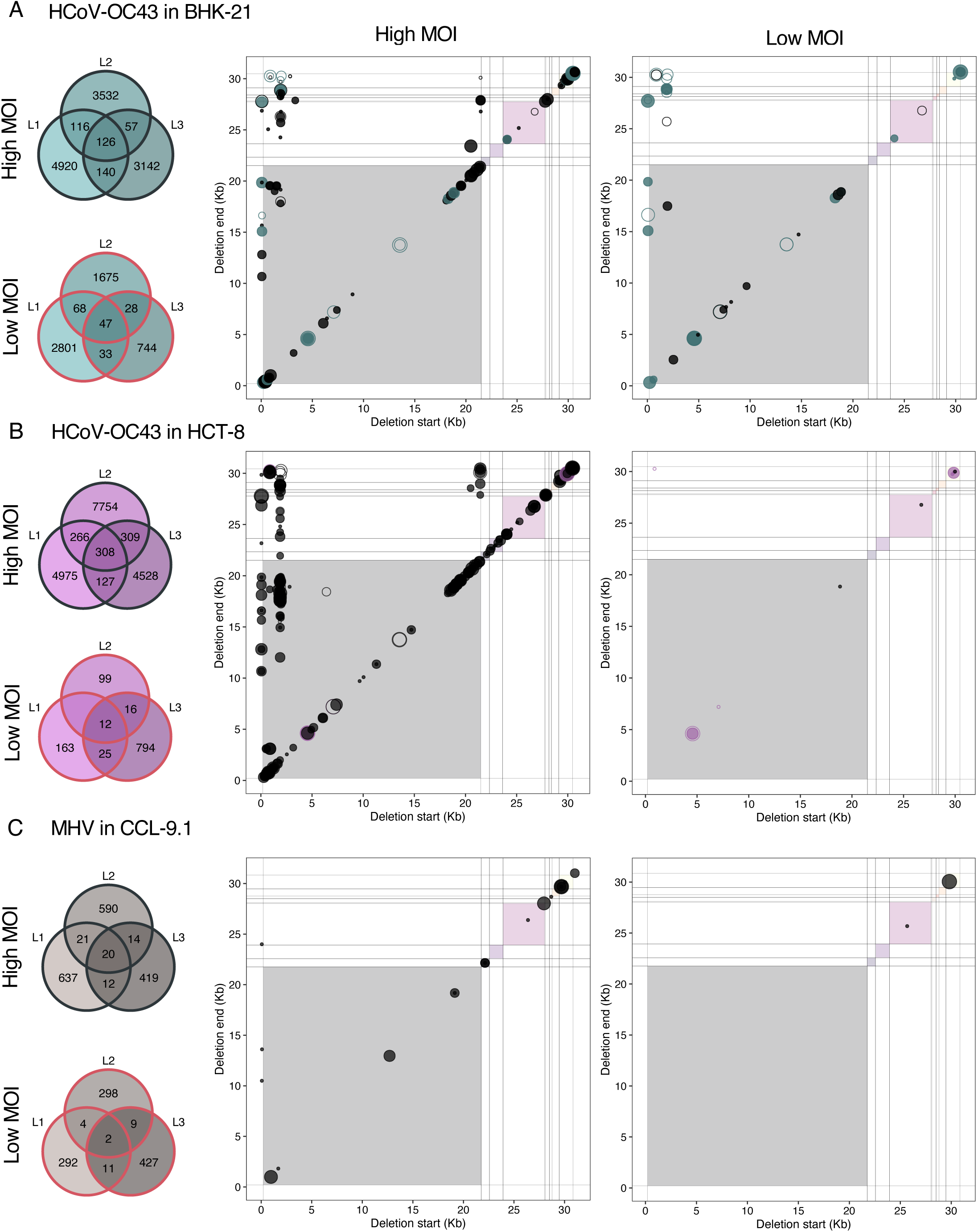
Convergencies between evolutionary lineages. Intersection size of deletions founded at lineages of HCoV-OC43 evolved in BHK-21 at **(A)** high and **(B)** low MOIs. BP and RP coordinates of common deletions shared by all the HCoV-OC43 lineages evolved in BHK-21cells at **(C)** high and **(D)** low MOIs. Intersection size of deletions founded at lineages of HCoV-OC43 evolved in HCT-8 at **(E)** high and **(F)** low MOIs. BP and RP coordinates of common deletions of parallel lineages of HCoV-OC43 in HCT-8 at **(G)** high and **(H)** low MOIs. Intersection size of deletions founded at lineages of MHV evolved in CCL-9.1 at **(I)** high and **(J)** low MOIs. For all the scatter plots, the empty points indicate the deletion was already found in the ancestral virus; if the dot color is different form black, the deletion was convergent also between both MOIs. Background colors represent cistrons as in Fig. 8A for HCoV-OC43 and Fig. 8B for MHV. Symbols’ size indicates the number of samples where the deletions were found.

Noticeably, despite their scarceness, for both viruses and regardless the cell types the count of shared DVGs among the three lineages consistently remained higher at high MOI (126 *vs* 47 and 308 *vs* 12, for HCoV-OC43 in BHK-21 and HCT-8, respectively; and 20 *vs* 2 for MHV).

Subsequently, to further characterize the few common deletions shared by the three parallel lineages, we examined their coordinates. In HCoV-OC43 samples (Fig. 9A), the coordinates of deletion BP and RP, paralleled with the coordinates of the persistent deletions shown in Fig. 8B. Again, we observed that the most frequent breakpoints were at the 5’ end of the genome for both cell types and MOIs (within the first 2 kb) and in the last third part of the 5’ genome (around 21 kb) appeared at high MOI, here more specifically in the HCT-8 samples. Similar to the case of persistent DVGs, we observed a limited number of deletions BP but highly variable RP.

Finally, we examined the impact of cell type on the generation of common deletions among HCoV-OC43 lineages (Fig. 10). Consistent with the probabilistic computations detailed above, we observed a substantial deviation between the actual counts in each of the three categories and the expected count distribution for both MOIs (χ² ≥ 28,669.853, 2 d.f., *P* < 0.001). This effect primarily steamed from a shortage of shared DVGs between the two cell types, suggesting that the spectrum of DVGs generated by HCoV-OC43 was influenced, to some extent, by the specific cell in which it was replicating. Notably, this deficit was more pronounced for lineages evolved at high MOI. Additionally, when comparing the count numbers in the three groups for lineages evolved at high and low MOIs, we discovered a highly significant distinction (homogeneity χ² = 4228.490, 2 d.f., *P* < 0.001). Specifically, at high MOI, lineages evolved in HCT-8 cells contained proportionally more unique DVGs than lineages evolved in BHK-21 cells, whereas the situation was reversed at low MOI (Fig. 10A).

**Figure 10.**
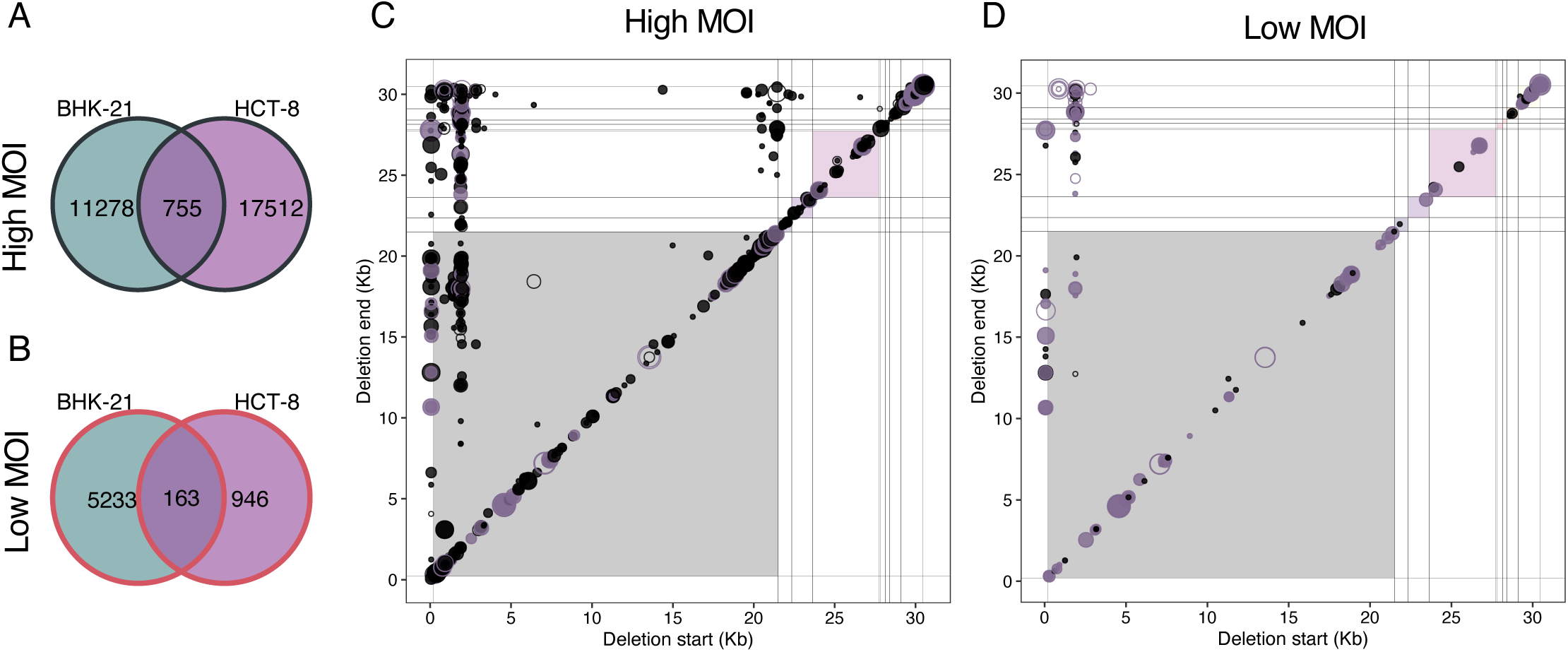
HCoV-OC43 deletion DVGs convergent among both cell types. (A-B) Intersection sizes at **(A)** high and **(B)** low MOIs. Hotspots of common deletions among cell types (intersection set in panels A and B) at **(C)** high and **(D)** low MOIs. Background colors indicate viral cistrons (as in Fig. 8A), the size of the points indicate the number of samples where the deletion is present. Empty circles indicate that the deletions were already present in the ancestral stock and purple and black colors represent, respectively, common to both MOIs or unique to the MOI.

It was noteworthy that 114 out of the 163 convergent deletions identified under low MOI conditions were also present in the high MOI set. More strictly, 60 deletions were shared across all lineages in both cell types at high MOI and six at low MOI. Additionally, the pattern of deletion coordinates shared between the two host cell types (Fig. 10C and 10D) was similar to the pattern of common deletions among parallel lineages of the same cell type (Fig. 9A) and also to the patterns of persistent deletions (Fig. 8B). This consistency could be taken as further evidence that not all deletions were random occurrences, but were regulated by similar molecular mechanisms or selective pressures.

## 3. Discussion

During replication, viruses, especially RNA viruses, spontaneously generate varying versions of their genomes, including hypermutated genomes, insertions, deletions and reorganizations (5’ and 3’ cb and sb). These unusual genomes cannot complete a whole viral cycle on their own but can play crucial roles in infection outcomes and viral evolution (Rezelj, Levi and Vignuzzi 2018; Vignuzzi and López 2019; Aguilar Rangel et al. 2023; Elena 2023). Understanding the dynamics of DVG formation and accumulation can provide insights into the underlying mechanisms of their appearance and function. To investigate this in betacoronaviruses, we conducted evolution experiments with HCoV-OC43 and MHV.

DVGs exhibited an overall increase in both richness and abundance in all evolving lineages. In HCoV-OC43, DVG richness initially spiked out but later stabilized, suggesting the existence of a diversity ceiling. However, DVG abundance at high MOI remained consistently high in advanced passages. At high MOI, HCoV-OC43 DVGs showed higher variability and abundance compared to low MOI, likely due to potential *trans*-complementation and other sort of interactions between DVGs and helper viruses. Our observations support the hypothesis that at high MOI, DVGs in HCoV-OC43 were not only generated *de novo* after each passage but some might also be selected, favoring the accumulation of the fittest DVGs while leading to a decrease in less fit variants. Indeed, at high MOI, much larger proportion of persistent DVGs was observed compared to low MOI. Transmission at low MOI likely occurred through secondary infections, as uninfected cells remained available after the first round of infection. Additionally, transmission could be facilitated by collective viral infection (Maeda et al. 1999; Cuevas, Durán-Moreno and Sanjuán 2017; Andreu-Moreno and Sanjuán 2018; Rosell et al. 2021; Zavan et al. 2023) or cell fusion (Buchrieser et al. 2020; Leroy et al. 2020; Rajah et al. 2022; Zavan et al. 2023). Under these conditions, a DVG could co-infect with a helper virus and be *trans*-complemented for successful propagation, even at an initially low MOI, despite of foundational effects could be more accused at low MOI and in some of the lineages the viral load got undetectable.

Significant DVG accumulation was observed in both cell lines with HCoV-OC43, in BHK-21 cells there was also a general increase with small fluctuations in infectious viral load. The positive trend for both full and defective virus suggest that not necessarily an interfering dynamic was occurring. These dynamics reinforce the idea of DVGs as triggers and/or reservoirs of viral diversity, providing an adaptative advantage (Vignuzzi and López, 2019; Gribble et al. 2021; Lin et al. 2023). However, in HCT-8 cells, viral loads showed an overall decreasing trend. This could be attributed to technical limitations in testing viral loads, as it had to be assessed in BHK-21 cells due to the inability to form plaques in HCT-8 cells. Viral adaptation to HCT-8 cells during the passages might have resulted in fitness tradeoffs in the BHK-21 cells used for plaquing. Further studies are needed to investigate the potential role of DVG - complete virus cooperation at this stage.

In contrast to HCoV-OC43, the richness and abundance of DVGs in MHV were positively correlated and increased throughout evolution, while the overall trend for viral load indicated a decrease, leading to undetectable levels in some MOI lineages. We proposed two non-mutually exclusive hypotheses to explain the observed fluctuations in viral load, one calling for the increasing accumulation of DVGs and another one associating the effect to reductions in the inoculum size and the subsequent onset of Muller’s ratchet (Clarke et al. 1993). Our finding of an accumulation of diverse populations of DVGs in evolving lineages, in both viruses but particularly in MHV, gives support to the first possibility, although does not rule out the second hypothesis, specially at low MOI.

The MHV strain that we used in this experiment was interferon-sensitive. Although MHV has a short infection cycle and should not be notably affected by interferon-based cell defense (Kyuwa and Sugiura 2020), we cannot completely rule out the possibility that interplay between the virus and the cells’ immune response may have impacted the generation and propagation of DVGs, contributing too to the divergent outcomes observed amongst the two betacoronaviruses. The interferon response has been identified as key in infections in which the presence of DVGs impacts the severity of symptoms (Zhou et al, 2023), or in which infection does not resolve but becomes chronic (Mura et al. 2017; Vignuzzi and López 2019; Wignall-Fleming et al. 2020).

The choice of cell lines for the HCoV-OC43 evolution experiments was primarily motivated by their high susceptibility and ability to produce high viral titers, which was essential for virus passaging at a high MOI. As a result, the outcome of viral DVG evolution in both cell lines showed similarities, indicating that they may not have imposed largely different constraints for viral adaptation. In retrospective, the use of more diverse cell lines with varying properties and susceptibilities might had provided additional insights into the impact of host factors on DVG evolution. Nevertheless, the results obtained from the selected cell lines still offered valuable information on DVG dynamics and accumulation in the context of HCoV-OC43 evolution. Future studies could consider employing a broader range of cell lines to gain a more comprehensive understanding of how host factors influence the evolution of DVGs. Indeed, Aguilar Rangel et al. (2003) have shown that Dengue virus (DENV) generates different spectra of DVGs in human and mosquito cells. Likewise, Hasiów-Jaroszewska et al. (2018) also described variation in the composition of DVGs of tomato black ring virus in different host species. We could also compare our results for MHV in CCL-9.1 with those obtained in DBT-9 cells by Gribble et al. (2021) and identify common hotspots of recombination, as well as differences.

Deletions were the most variable and abundant type of reconstructed DVGs, an observation that is consistent with previous studies with other positive-sense single-stranded RNA viruses: Jaworski and Routh (2017) for Flock House virus, Aguilar Rangel et al. (2023) in the cases of poliovirus and DENV, and Zhou et al. (2023) in the case of SARS-CoV-2. For this reason, we conducted a more detailed analysis of this specific DVG type. The analysis, focusing on deletions longer than three nucleotides, distinguish patterns in genome sizes that dominated in abundance. Over the course of HCoV-OC43 evolution at high MOI, these patterns refined within a specific size range, giving rise to dynamic peaks of high abundance. This dynamic behavior within a certain size range suggests adaptive evolution. In contrast, the accumulation pattern of certain sizes at low MOI was not prominent, and, notably, MHV did not exhibit a clear pattern of size accumulation. This lack of a discernible pattern may indicate different evolutionary dynamics or constraints under low MOI. Furthermore, short genomes with deletions (< 2 kb) were particularly prevalent in the HCT-8 cell lineage. This observation underscores the potential influence of host-specific factors on the evolution dynamics of viral genomes with deletions.

Furthermore, we observed that the cistrons E and ns12.9 (or its syntenic accessory protein 4 in MHV) were the more impacted by deletions, while ORF1ab and S were the less affected at both MOIs for both viruses. Coronaviruses E cistron encodes for a small structural protein involved in multiple stages of the replicative cycle, it participates in viral assembly, virions release and pathogenesis (Schoeman and Fielding 2019). In MHV, E is not essential for virus replication and a ΔE mutant showed a lower growth rate and a lower infectious titer than the wildtype virus, but it still was capable of producing viable particles (Kuo and Masters 2003). E is highly expressed in infected cells, but only a small part is incorporated to the virion envelope. E also can form homotypic interactions, oligomerizing to generate viroporins, ion-channel proteins associated with pathogenicity. HCoV-OC43 ns12.9 protein is associated with virus morphogenesis and pathogenesis by viroporin formation. The HCoV-OC43-Δns12.was defective in growth *in vitro* and a reduced inflammation and virulence in the brain of infected mice, confirming its influence in virus pathogenesis (Zhang et al. 2015). Some studies had associated the inhibition of the E viroporins with a reduction of the pathogenicity, highlighting its therapeutical value to face coronavirus infections (Schoeman and Fielding 2019), while others point towards different viroporins as potential targets for antiviral drugs (Zhang et al. 2015). The identification of E, ns12.9 and accessory protein 4 as the most prone cistrons for deletions suggests the tantalizing possibility that DVGs in these cistrons may be selected for as a viral strategy to modulate virulence.

Furthermore, the E protein may interact with the host’s immune system, implying that deletions in these regions could impact the virus ability to replicate and evade immune responses. Our evolution experiment took place in cell culture and not in a whole organism with complete immune system. The independent emergence of deletions of equivalent parts of genome in different viruses and cell types suggests that the shift from the whole organism to cell culture was a more substantial landmark than the differences among various cell culture types. Additionally, these deletions were accumulated already in the first stages of the evolution, which suggests that the adaptive advantage of these variants was significant. However, the underlying mechanisms could be much more complex, potentially involving specific viral sequences and structures. The analysis of the most frequent sites for BP and RP, along with their surrounding sequences, suggests that the mechanism of DVG generation is not random and is influenced, in part, by the genome sequence but further by the resulting secondary RNA structure.

It is noteworthy that some hotspots of persistent deletions BP are in the first third of the viral genome, a region generally assumed to be crucial for viral replication. We identify a site close to the TRS-L as one of the most frequent BP, suggesting that the RNA structures needed for replication would be intact. We should not rule out the effect that successive deletions have on the increase in frequency in some spots, or a special case of the latter, that sgRNAs are targets of deletions. Unfortunately, the sequencing technology used for this work limits our ability to investigate this further.

We have also paid attention to identify deletions that consistently appeared over the course of evolution. In the case of HCoV-OC43, some of these persistent deletions likely represent instances where deletions appeared *de novo* multiple times in the same lineage, because our results also indicated that the process of DVG generation was not completely random. Notably, the pattern of coordinates for persistent deletions highlighted the existence of convergent deletions between parallel lineages and even between different cell types. This suggests that a subset of deletions arose independently of the host and lineage. Another portion of pervasive but not *de novo* generated deletions, was likely composed of deletions that were transmitted along passages. When comparing samples at high *vs* low MOIs, we observed more persistent deletions under high MOI conditions, suggesting that the virus, given the opportunity for *trans*-complementation, exhibits an increased tendency for supporting persistent deletions. In the case of MHV, most DVGs were not transmitted but *de novo* generated within each passage, not being under specific selection pressure, consistent with the less defined and more variable size distribution. This suggests, in general, that the dynamics of DVG accumulation differs among two closely related betacoronavirus species, with potentially distinct mechanisms governing their appearance, abundance and fitness (de Groot, van der Most and Spaan 1992; Brian and Spaan 1997).

Furthermore, our findings indicated that the dynamics of persistent deletions in HCoV-OC43 at high MOI maintained hotspots of deletion coordinates, becoming more abundant with passages. Certain genomic regions seemed more susceptible to deletions than others (Lin and Lai 1993), with these hotspots showing increased abundance during experimental evolution at high MOI. In both viruses, regions susceptible to deletions were detected, and the abundance of deletions in these hotspots increased with the progress of experimental evolution at high MOI. Despite a larger genome size being a disadvantage for replication competition, longer DVGs lacking structural proteins may have the advantage of being more readily complemented in *trans* and transmitted. Zhou et al. (2023) also found hotspots of BP and RP in the genome of SARS-CoV-2 in samples from patients, which suggest that our *in vitro* results might reflect relevant processes *in vivo*.

The dynamics of deletions in parallel lineages showed a remarkable degree of convergence, despite the majority of shared deletions between the lineages were very short, potentially classified as sequencing errors. After removing these short ones, we still found a number of DVGs of various sizes shared among lineages evolved in the same host. This finding prompted for distinct patterns of deletion evolution within different cell types, combining stochastic generation of deletions within each passage and the potential persistence of positively selected deletions shared by different lineages. The observation that some deletions can be positively selected to modulate virus replication and pathogenesis, for instance by altering coding reading frames and generating novel proteins (Aguilar Rangel et al. 2023), opens tantalizing possibilities for the development of DVGs with antiviral activity. In general, short DIPs, and not long DVG, have been considered for their interfering activity over wild-type viruses (Notton et al. 2014), although results by Levi et al. (2021) suggest that long DVGs can be effective and broad-spectrum antiviral for alphaviruses.

In conclusion, the buildup of DVGs seems to be governed by dynamic processes that set a cap on the diversity of DVGs capable of accumulating in each sample. This restriction is affected by both the passage number and, in specific instances, the MOI. Throughout evolution at high MOI, a selective process appears to take place, giving preference to certain DVGs for accumulation while others diminish over successive passages. This observation implies that DVGs possess the ability to transmit under high MOI conditions, resulting in their enduring presence and potential influence on viral evolution.

## 4. Material and Methods

### 4.1. Viruses and cells

Human large intestine carcinoma cells (HCT-8; ATCC^®^ CCL-244^TM^) and murine liver cells NCTC 1469 (CCL-9.1; ATCC^®^ CCL-9.1^TM^) were obtained from the American Type Culture Collection (ATCC). Baby hamster kidney cells (BHK-21) were kindly provided by Dr. R. Sanjuán (I^2^SysBio, CSIC-UV). BHK-21 cells and HCT-8 cells were cultured at 37 °C in DMEM (Gibco-Invitrogen) supplemented with 0.22% (w/v) sodium bicarbonate (Sigma Aldrich), sodium pyruvate (Sigma Aldrich), 10% fetal bovine serum (FBS), 1× penicillin-streptomycin (Gibco-Invitrogen), 1× amphotericin B (Gibco-Invitrogen), and non-essential amino acids using standard laboratory procedures. CCL-9.1 cells were cultured at 37 °C in DMEM media supplemented with 0.22% (w/v) sodium bicarbonate (Sigma Aldrich), sodium pyruvate (Sigma Aldrich), 10% horse serum, 1× penicillin-streptomycin, 1× amphotericin B, and non-essential amino acids. All cell lines were routinely tested for mycoplasma contamination.

The viruses used in this study were HCoV-OC43 (ATCC^®^ VR-1558) and MHV (ATCC^®^ VR-766), both obtained from ATCC. The maintenance medium for HCoV-OC43 was DMEM supplemented with 0.22% (w/v) sodium bicarbonate (Sigma Aldrich), sodium pyruvate (Sigma Aldrich), 2% FBS, 1× penicillin-streptomycin (Gibco-Invitrogen), and 1× amphotericin B (Gibco-Invitrogen). Cell culture media of CCL-9.1 was also used as maintenance medium by infection with MHV.

### 4.2. Experimental evolution

To generate the initial virus inoculum, BHK-21 and HCT-8 cells were plated one day before and growth to 70 - 80% confluency. Afterward, the cell layers were infected with the corresponding virus at a MOI of 0.01. HCoV-OC43 was grown for 4 d at 33 °C, and MHV was grown for 20 h at 37 °C. The supernatants from infected cells were then collected and used as the infectious stock. For successive passages, all cell types were grown in 6-well plates to 70 - 80% confluency and infected with 250 µL of undiluted viral inoculum from the previous passage. HCoV-OC43 was passaged 47 times in BHK-21 cells and 31 times in HCT-8 cells, while MHV was passaged 19 times in CCL-9.1 cells.

The MOI was not maintained the same throughout all passages; instead, at high MOI, we infected every passage without dilution to get highest MOI possible. For low MOI samples, the inocula were diluted to avoid infecting each cell with more than one infectious particle. Supplementary Table S1 shows the range of MOIs for each virus and cell type.

### 4.3. Viral load quantification

The quantity of infectious virus in the supernatant was determined after each passage. To determine the infectious viral particles for HCoV-OC43, plaque assays were performed in BHK-21. The procedure involved seeding the cells into 6-well plates and incubating them for 24 h until reaching 80% confluency. Serial dilutions of the virus were prepared in DMEM media, and 250 μL of each dilution was used to infect the wells of the 6-well plate for 90 min at 33 °C and 5% CO_2_ with swinging every 15 min. A negative control consisting of only DMEM media without virus was included for comparison. After the infection period, the cells were covered with 2 mL of media, which consisted of a 50:50 mix of 2× DMEM medium supplemented with 2% FBS and 2% agar. The agar-DMEM layer was overlaid with liquid DMEM medium supplemented with 1% FBS as it solidified. The plates were then incubated at 33 °C and 5% CO_2_ for 5 d. After incubation, cells were fixed using 10 % formaldehyde and plugs were removed. Monolayers were stained with 2 % crystal violet in 10 % formaldehyde, washed with tap water, and plaques counted to determine the plaque forming units (PFU) per mL of inoculum.

The infectious viral particles for MHV were determined by TCID_50_. Cells were seeded into 96-well plates and incubated for 24 h to reach 90% confluence. Viral transfer samples were diluted in DMEM complete media. Once diluted, 125 μL of the dilutions for each sample were used to infect the wells of the 96-well plate. A negative control consisting of only DMEM complete media was also included. The plates were then incubated at 37 °C for 2 d. After incubation, the wells were inspected under a light microscope for the presence of visual cytopathic effect, and the TCID_50_/mL was calculated using the Reed and Muench (1938) method. For comparison purposes with plaque assay results, TCID_50_/mL values were converted into PFU/mL by multiplying by ln(0.5) ý 0.693.

### 4.4. Preparation of viral RNA

Samples from four equidistant time series of each evolutionary lineage were collected to extract RNA using the NZY Viral RNA Isolation kit (NZYTech) following the manufacturer’s instructions. After extraction, the RNA was purified and concentrated using the RNA Clean&Concentrator-5 Kit (Zymo Research).

### 4.5. High-throughput RNA sequencing

Libraries for high-throughput genome sequencing (HTS) were prepared by Novogene UK Company Ltd (www.novogene.com) using at least 200 ng of total RNA or a larger amount if available. The ribosomal RNAs (rRNA) from both eukaryotes and prokaryotes were depleted from the total RNA samples. The remaining RNAs were fragmented into 250 - 300 bp fragments and reverse-transcribed into double-stranded cDNAs, followed by end repair, A tailing, and adapter ligation. After fragment size selection and PCR amplification, the metatranscriptome library was checked for quality, and sequencing was conducted on the Illumina platform. The raw data obtained from the samples ranged between 3 Gb and 15 GB, providing ∼64 - 124,981 ×viral nucleotide mean depth, with a median of 13,362.52. The sequencing was not strand-specific.

### 4.6. Processing of HTS reads and identification of DVGs

Raw paired reads from Illumina metagenomic data from each sample were merged into a single interleaved file using the reformat.sh script from the BBMap/BBTool package (https://github.com/BioInfoTools/BBMap/blob/master/sh/reformat.sh). The interleaved reads were then processed using the BBduk.sh tool to remove adapter sequences, clean, and trim the reads to obtain high-quality nucleotides at both ends (https://github.com/BioInfoTools/BBMap/blob/master/sh/bbduk.sh). Next, the remaining reads were mapped to the corresponding host genomes: the hamster genome for BHK-21 cell lineages, the human genome for HCT-8, and the mouse genome for CCL-9.1, using the BWA-MEM algorithm (Li 2013). Reads that mapped with the host genomes were excluded from further analysis. The resulting reads were analyzed using the metasearch tool DVGfinder to identify and classify DVG events. Predicted DVG species sizes were calculated based on the BP and RP by the program (Olmo-Uceda et al. 2022). Reference genomes used were ATCC VR-1558 for HCoV-OC43 and GU593319.1 for MHV.

The raw sequencing data were deposited in GenBank (NCBI) under BioProject number PRJNA1021788.

### 4.7. Processing and diversity measures of DVG

Only the results from DVGfinder Filtered mode were used. In the case where both possible DVG directions were present in the original results, the sum of their abundance values was used and BP and RP lost their original meaning. The canonic sgRNAs where identified and discarded from the rest of the analysis. A deletion-type DVG was identify as sgRNAs when its start coincides with the TRS-leader and its end with one of the canonical TRS-backs, both coordinates with a flexible margin of ten bases.

Richness was understood as the number of different DVGs relative to the total number of mapped viral reads of the group (sample or lineage, depending on the case) per 10^5^ (RPHT). Abundance was a used as a measure of the number of reads supporting the DVG, relative to the total number of mapped viral reads of the group (sample or lineage, depending on the case) per 10^5^.

### 4.8. Identification of transmitted DVGs

To study the possible transmission of DVGs between passages, each DVG was labeled as persistent if the exact same event (DVG type with same BP and RP coordinates) was identified in a previous passage of the same lineage (including the stock).

### 4.9. Characterization of deletion type DVGs

All the DVGs with deletions bigger than 3 bases were analyzed from different perspectives: their length, their location in the genome, their distance to the nearest TRS, their nucleotide composition (Fig. 5E) and their secondary structure (Fig. 5F). The cistrons more affected by deletions were also identified.

#### 4.9.1. Deletion size distribution

The distribution of the reconstructed size of deletion type DVG was calculated considering the abundance of each specie in the sample. For a better visualization of the frequency coordinates only deletions bigger than 50 bases were represented (Fig. 4A and Supplementary Fig. S2). The size of the deleted fragments was analyzed by a Lomb-Scargle periodogram method (VanderPlas 2018).

#### 4.9.2. Distribution of deletion start and ends in the viral genomes

For every cell type, a histogram of the start and end position of deletions were represented in Fig 5A. The data was generated with the method *hist* from the base package graphics setting the breaks argument to the quotient of the viral length between 50.

#### 4.9.3. *K*-mer analysis

The *k*-mer composition of the sequences ten bases pre- and post-start and end of deletions was compared with the *k*-mer composition of the complete genome (*k* = 1 to 3) with a χ² test (*chisq.test* from base package *stats*, with *rescale.p = T*). The *k*-mer composition was analyzed with the *oligonucleotideFrequency* method from Biostrings v2.66.0 (Pagès et al. 2023).

#### 4.9.4. Minimum Free Energy (MFE) around junctions

The MFE of the structures predicted with RNAfold v2.5.1 (Lorenz et al. 2016) for 50-base sliding windows were used to perform a Spearman correlation analysis with the frequencies for 50-base windows of being start or end coordinates of a deletion calculated previously. RNA2Drawer (Johnson et al. 2019) was used for representation of the obtained structures.

#### 4.9.5. Deletions per base calculation

To evaluate the viral cistrons more affected by deletions the *findOverlaps* method from GenomicRanges v1.50.2 (Lawrence et al. 2013) was used in a homemade script in R version 4.3.2. The number of deletions was relativized to the cistron length.

### 4.10. Other statistical analyses

Except otherwise indicated, statistical analyses were done with SPSS version 28.0.1.0 (IBM, Armonk NY, USA).

Viral load time series for individual lineages were fitted to the general ARIMA(*p*, *d*, *q*) model:

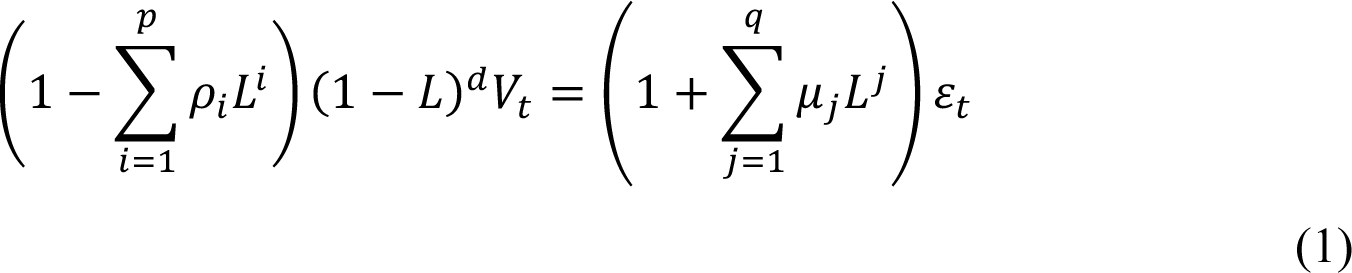

where *V_t_* represents the viral load at passage *t*, *L* is the lag operator such that *L^n^V_t_* = *V_t_* _−_*_n_*, *π_i_* are the parameters of the autoregressive part of the model with *p* := 0 being the order of the autoregressive model (number of time lags), *μ_i_* the parameters of the moving average part with *q* := 0 being the order of the moving-average model, *d* := 0 is the differencing order, representing the number of times the time series is differenced to achieve stationarity, and *ρ._t_* are the error terms assumed to be independent identically distributed *N*(0, *σ*^2^). ARIMA fitting and model selection was done using the package *forecast* in R version 4.3.2 in RStudio version 2023.12.1+402. Estimated slope values were further analyzed by runs tests or pairwise Mann-Whitney tests.

Diversity and abundance of DVGs classes were analyzed using the Scheirer-Ray-Hare (SRH) non-parametric two-ways analysis of variance. In all tests done, the specific factor analyzed (*i.e*., virus species, cell type or MOI) were orthogonal to the class of DVG (*i.e*., deletions, insertions, 3’ and 5’ cb).

Other specific statistical tests are presented in the text as needed.

### 4.11. Code implementation

All the code developed for this work is accessible in https://github.com/MJmaolu/AccumulationDynamicsDVGs/tree/main. Intensive computations were run on the HPC cluster Garnatxa at I2SysBio (CSIC-UV). The R scripts were run in Rstudio version 4.2.1 (2022-06-23).

## Supplementary data

Supplementary data are available at *Virus Evolution* on line.

## Acknowledgements

We thank Paula Agudo, Juan V. Bou, José L. Carrasco, Francisca de la Iglesia, J. Tomás Lázaro, María C. Marqués, Josep Sardanyés, and Ernesto Segredo-Otero for help, assistance and fruitful discussions. This work was supported by CSIC PTI Salud Global grant 202020E153, by grants SGL2021-03-009 and SGL2021-03-052 from European Union NextGenerationEU/PRTR through the CSIC Global Health Platform established by EU Council Regulation 2020/2094, and by grant PDC2022-133020-I00 from MCIN/AEI/10.13039/501100011033 and “European Union NextGenerationEU/PRTR”. M.J.O-U. was supported by grant FPU2019/05246 funded by MCIN/AEI/10.13039/501100011033 and “ESF investing in your future”.

## Supplementary materials

**Figure S1.** Percentage of sgRNAs over the total number of deletions observed per sample. Thick lines are only shown for illustrating the average trend over time.

**Figure S2.** Distribution of deletion type DVGs at low MOI. To facilitate visualization only deletions bigger than 50 nucleotides are shown. The number represent the exact length of the DVG and in parenthesis the lineages where they were founded.

## References

1. Aguilar Rangel, M., et al. (2023) ‘High-resolution mapping reveals the mechanism and contribution of genome insertions and deletions to RNA virus evolution’, Proceedings of the National Academy of the USA, 120:e2304667120.

2. Andreu-Moreno, I. and R. Sanjuán (2018) ‘Collective infection of cells by viral aggregates promotes early viral proliferation and reveals a cellular-level Allee effect’, Current Biology, 28:3212–3219.

3. Barrett, A. D. T. and N. J. Dimmock (1984) ‘Modulation of Semliki Forest virus-induced infection of mice by defective-interfering virus’, Journal of Infectious Diseases, 150:98–104.

4. Belshaw, R., R. Sanjuán and O. G. Pybus (2011) ‘Viral mutation and substitution: units and levels’, Current Opinion in Virology, 1:430–435.

5. Brian, D. A. and R. S. Baric (2005) ‘Coronavirus genome structure and replication’, Current Topics in Microbiology and Immunology, 287:1–30.

6. Brian, D. A. and W. J. M. Spaan (1997) ‘Recombination and coronavirus defective interfering RNAs’, Seminars in Virology, 8:101–111.

7. Buchrieser, J. et al. (2020) ‘Syncytia formation by SARS-CoV-2-infected cells’, EMBO Journal, 39:e106267.

8. Clarke, D. K. et al. (1993) ‘Genetic bottlenecks and population passages cause profound fitness differences in RNA viruses’, Journal of Virology, 67:222–228.

9. Cuevas, J., M. Durán-Moreno and R. Sanjuán (2017) ‘Multi-virion infectious units arise from free viral particles in an enveloped virus’, Nature Microbiology, 2:17078.

10. De Groot, R. J., R. G. van der Most and W. J. M. Spaan (1992) ‘The fitness of defective interfering murine coronavirus DI-a and its derivatives is decreased by nonsense and frameshift mutations’, Journal of Virology, 66:5898–5905.

11. De Haan, C. A. M. et al. (2002) ‘Coronaviruses maintain viability despite dramatic rearrangements of the strictly conserved genome organization’, Journal of Virology, 76:12491–12502.

12. De la Iglesia, F. and S. F. Elena (2008) ‘Fitness declines in tobacco etch virus upon serial bottleneck transfers’, Journal of Virology, 81:4941–4947.

13. Di Gioacchino, A. et al. (2022) ‘sgDI-tector: Defective interfering viral genome bioinformatics for detection of coronavirus subgenomic RNAs’, RNA, 28:277–289.

14. Duarte, E. A. et al. (1993) ‘Many-trillionfold amplification of single RNA virus particles fails to overcome the Muller’s ratchet effect’, Journal of Virology, 67:3620–3623.

15. Duffy, S., L. A. Shackelton and E. C. Holmes (2008) ‘Rates of evolutionary change in viruses: patterns and determinants’, Nature Reviews Genetics, 9:267–276.

16. Elena, S. F. (2023) ‘The role of indels in evolution and pathogenicity of RNA viruses’, Proceedings of the National Academy of Sciences of the USA, 120:e2310785120.

17. Elena, S. F. et al. (1996) ‘Evolution of fitness in experimental populations of vesicular stomatitis virus’, Genetics, 142:673–679.

18. Elshina, E., & te Velthuis, A. J. W. (2021). ‘The influenza virus RNA polymerase as an innate immune agonist and antagonist’. In Cellular and Molecular Life Sciences (Vol. 78, Issue 23, pp. 7237–7256). Springer Science and Business Media Deutschland GmbH.

19. Felt, S. A. et al. (2022) ‘Accumulation of copy-back viral genomes during respiratory syncytial virus infection is preceded by diversification of the copy-back viral genome population followed by selection’, Virus Evolution, 8:veac091.

20. Fosmire, J. A., K. Hwang and S. Makino (1992) ‘Identification and characterization of a coronavirus packaging signal’, Journal of Virology, 66:3522–3530.

21. Goebel, S. J., J. Taylor and P. S. Masters (2004) ‘The 3’ *cis*-acting genomic replication element of the severe acute respiratory syndrome coronavirus can function in the murine coronavirus genome’, Journal of Virology, 78:7846–7851.

22. Gribble, J. et al. (2021). ‘The coronavirus proofreading exoribonuclease mediates extensive viral recombination’, PLoS Pathogens, 17:e1009226.

23. Hasiów-Jaroszewska, B. et al. (2018) ‘Defective RNA particles derived from tomato black ring virus genome interfere with the replication of parental virus’, Virus Research, 250:87–94.

24. Hofmann, M. A., P. B. Sethna and D. A. Brian (1990) ‘Bovine coronavirus mRNA replication continues throughout persistent infection in cell culture’, Journal of Virology 64:4108–4114.

25. Huang, A. S. (1973) ‘Defective interfering viruses’, Annual Review of Microbiology, 27:101–118.

26. Huang, A. S. and D. Baltimore (1970) ‘Defective viral particles and viral disease processes’, Nature, 226:325–327.

27. Jaworski, E. and A. Routh (2017) ‘Parallel ClickSeq and nanopore sequencing elucidates the rapid evolution of defective-interfering RNAs in Flock House virus’, PLoS Pathogens, 13:e1006365.

28. Jennings, P. A. et al. (1983) ‘Does the higher order structure of the influenza virus ribonucleoprotein guide sequence rearrangements in influenza viral RNA?’, Cell, 34:619–627.

29. Johnson, P. Z. et al. (2019) ‘RNA2Drawer: geometrically strict drawing of nucleic acid structures with graphical structure editing and highlighting of complementary subsequences’, RNA Biology, 16:1667–1671.

30. Kim, D. et al. (2020) ‘The architecture of SARS-CoV-2 transcriptome’, Cell, 181: 914–921.

31. Kuo, L. and P. S. Masters (2003) ‘The small envelope protein E is not essential for murine coronavirus replication’, Journal of Virology, 77:4597–4608.

32. Kyuwa, S., and Y. Sugiura (2020) ‘Role of cytotoxic T lymphocytes and interferon-ψ in coronavirus infection: lessons from murine coronavirus infections in mice’, Journal of Veterinary Medical Sciences, 82:1410–1414.

33. Lawrence, M. et al. (2013) ‘Software for computing and annotating genomic ranges’, PLoS Computational Biology, 9:e1003118.

34. Leroy, H. et al. (2020) ‘Virus-mediated cell-cell fusion’, International Journal of Molecular Sciences, 21:9644.

35. Levi, L. I. et al. (2021) ‘Defective viral genomes from chikungunya virus are broad-spectrum antivirals and prevent virus dissemination in mosquitoes’, PLoS Pathogens, 17:e1009110.

36. Li, T. and A. K. Pattnaik (1997) ‘Replication signals in the genome of vesicular stomatitis virus and its defective interfering particles: identification of a sequence element that enhances DI RNA replication’, Virology, 232:248–259.

37. Li, H. (2013) ‘Aligning sequence reads, clone sequences and assembly contigs with BWA-MEM’, *arXiv*, http://arxiv.org/abs/1303.3997.

38. Lin, Y. J. and M. M. Lai (1993) ‘Deletion mapping of a mouse hepatitis virus defective interfering RNA reveals the requirement of an internal and discontiguous sequence for replication’, Journal of Virology, 67:6110–6118.

39. Lin, C. H. et al. (2023) ‘Evolution of the coronavirus spike protein in the full-length genome and defective viral genome under diverse selection pressures’, Journal of General Virology, 104:001920.

40. Liu, P. et al. (2007) ‘A U-turn motif-containing stem-loop in the coronavirus 5’ untranslated region plays a functional role in replication’, RNA, 13:763–780.

41. Liu, Q., R. F. Johnson and J. L. Leibowitz (2001) ‘Secondary structural elements within the 3’ untranslated region of mouse hepatitis virus strain JHM genomic RNA’, Journal of Virology, 75:12105–12113.

42. Lorenz, R., I. L. Hofacker and P. F. Stadler (2016) ‘RNA folding with hard and soft constraints’, Algorithms for Molecular Biology, 11:8.

43. Maeda, J., A. Maeda, and S. Makino (1999) ‘Release of coronavirus E protein in membrane vesicles from virus-infected cells and E protein-expressing cells’, Virology, 263:265–272.

44. Mura, M. et al. (2017) ‘Nonencapsidated 5’ copy-back defective interfering genomes produced by recombinant measles viruses are recognized by RIG-I and LGP2 but not MDA5’, Journal of Virology, 91:e00643–17.

45. Notton, T. et al. (2014) ‘The case of transmissible antivirals to control population-wide infectious disease’, Trends in Biotechnology, 32:400–405.

46. Novella, I. S. et al. (1995) ‘Size of genetic bottlenecks leading to virus fitness loss is determined by mean initial population fitness’, Journal of Virology, 69:2869–2872.

47. Olmo-Uceda, M. J. et al. (2022) ‘DVGfinder: a metasearch tool for identifying defective viral genomes in RNA-Seq data’, Viruses, 14:1114.

48. Pagès, H. et al. (2023) ‘Biostrings: Efficient manipulation of biological strings’, R package version 2.70.1, https://bioconductor.org/packages/Biostrings.

49. Poirier, E. Z. et al. (2016) ‘Low-fidelity polymerases of alphaviruses recombine at higher rates to overproduce defective interfering particles’, Journal of Virology, 90:2446–2454.

50. Rajah, M. M. et al. (2022) ‘The mechanism and consequences of SARS-CoV-2 spike-mediated fusion and syncytia formation’, Journal of Molecular Biology, 434:167280.

51. Reed, L. J. and H. Muench (1938) ‘A simple method of estimating fifty per cent endpoints’, American Journal of Epidemiology, 27:493–497.

52. Repass, J. F. and S. Makino (1998) ‘Importance of the positive-strand RNA secondary structure of a murine coronavirus defective interfering RNA internal replication signal in positive-strand RNA synthesis’, Journal of Virology, 72:7926–7933.

53. Rezelj, V. V., L. I. Levi and M. Vignuzzi (2018) ‘The defective component of viral populations’, Current Opinion in Virology, 33:74–80.

54. Rosell, A. et al. (2021) ‘Patients with COVID-19 have elevated levels of circulating extracellular vesicle tissue factor activity that is associated with severity and mortality’, *Arteriosclerosis*, Thrombosis, and Vascular Biology, 41:878–882.

55. Roux, L., A. E. Simon and J. J. Holland (1991) ‘Effects of defective interfering viruses on virus replication and pathogenesis *in vitro* and *in vivo*’, Advances in Virus Research, 40:181–211.

56. Saira, K. et al. (2013) ‘Sequence analysis of *in vivo* defective interfering-like RNA of influenza a H1N1 pandemic virus’, Journal of Virology, 87:8064–8074.

57. Sanjuán, R. et al. (2010) ‘Viral mutation rates’, Journal of Virology, 84:9733–9748.

58. Schoeman, D. and B. C. Fielding (2019) ‘Coronavirus envelope protein: Current knowledge’, Virology Journal, 16:69.

59. Sethna, P. B., S. L. Hung and D. A. Brian (1989) ‘Coronavirus subgenomic minus-strand RNAs and the potential for mRNA replicons’, Proceedings of the National Academy of Sciences of the USA, 86:5626–5630.

60. Smith, C. M. et al. (2016) ‘Defective interfering influenza RNA inhibits infectious influenza virus replication in human respiratory tract cells: a potential new human antiviral’, Viruses, 8:237.

61. Stampfer, M., D. Baltimore and A. S. Huang (1971) ‘Absence of interference during high-multiplicity infection by clonally purified vesicular stomatitis virus’, Journal of Virology, 7:409–411.

62. Sun, Y. et al. (2019) ‘A specific sequence in the genome of respiratory syncytial virus regulates the generation of copy-back defective viral genomes’, PLoS Pathogens, 15:e1007707.

63. Van der Most, R. G., P. J. Bredenbeek and W. J. M. Spaan (1991) ‘Domain at the 3’ end of the polymerase gene is essential for encapsidation of coronavirus defective interfering RNAs’, Journal of Virology, 65:3219–3226.

64. VanderPlas, J.T. (2018) ‘Understanding the Lomb-Scargle periodogram’, Astrophysical Journal Supplement Series, 236:16.

65. Vignuzzi, M., and C. López (2019) ‘Defective viral genomes are key drivers of the virus-host interaction’, Nature Microbiology, 4:1075–1087.

66. Wignall-Fleming, E. B. et al. (2020) ‘Innate intracellular antiviral responses restrict the amplification of defective virus genomes of parainfluenza virus 5’, Journal of Virology, 94:00246–20.

67. Yang, D. and J. L. Leibowitz (2015) ‘The structure and functions of coronavirus genomic 3’ and 5’ ends’, Virus Research, 206:120–133.

68. Zavan, B. et al. (2023) ‘Extracellular vesicles and viruses: two intertwined entities’, International Journal of Molecular Sciences, 24:1036.

69. Zhang, R. et al. (2015) ‘The ns12.9 accessory protein of human coronavirus OC43 is a viroporin involved in virion morphogenesis and pathogenesis’, Journal of Virology, 89:11383–11395.

70. Zhao, X., K. Shaw and D. Cavanagh (1993) ‘Presence of subgenomic mRNAs in virions of coronavirus IBV’, Virology, 196:172–178.

71. Zhou, T. et al. (2023) ‘Generation and functional analysis of defective viral genomes during SARS-CoV-2 infection’, mBio, 14:e00250–23.

72. Zuker, M. (1989) ‘On finding all suboptimal foldings of an RNA molecule’, Science, 244:48–52.

