## Supplementary figures and images for "Accumulation dynamics of defective genomes during experimental evolution of two betacoronaviruses"

### Supplementary Fig. S1

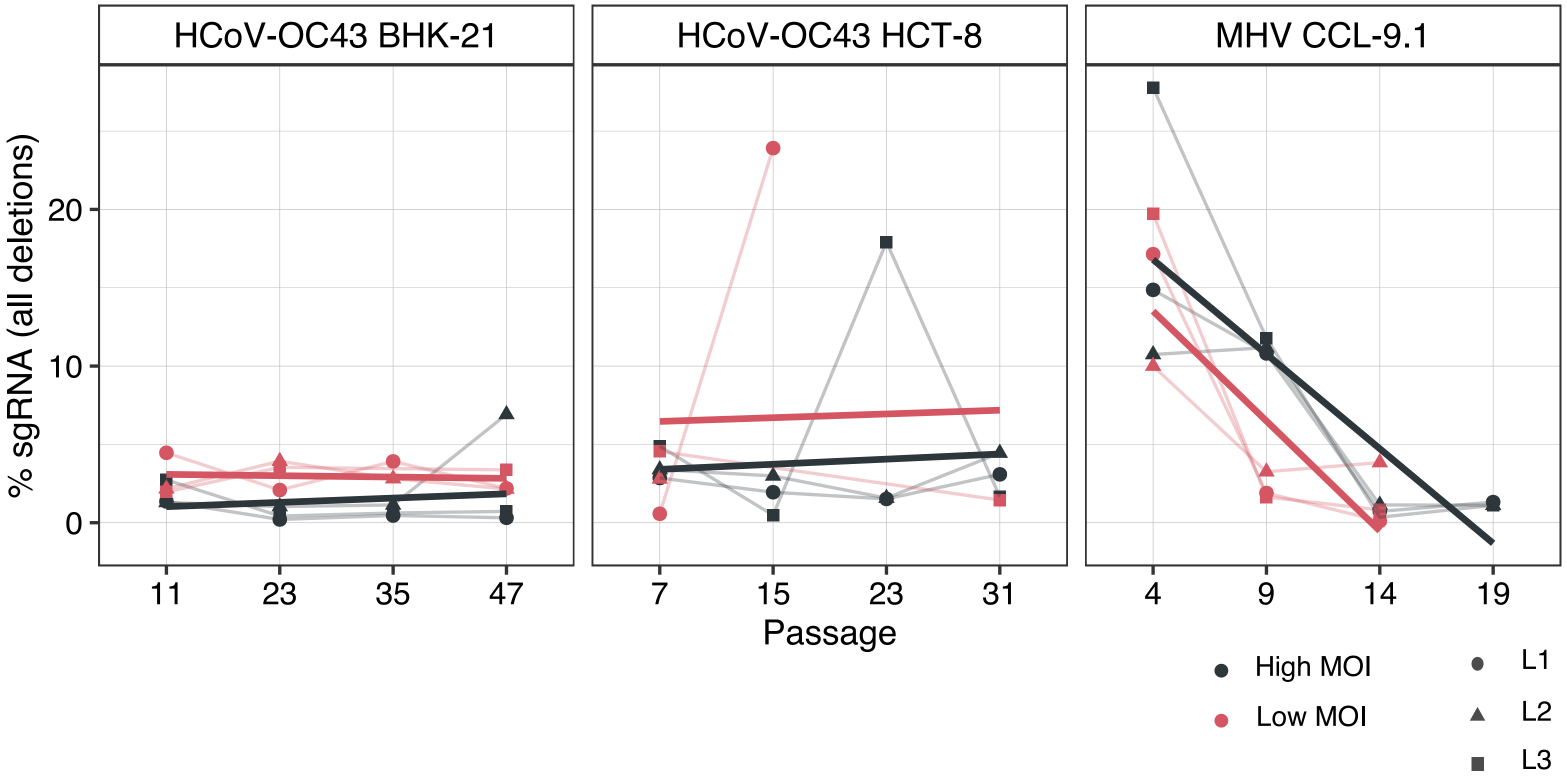

### Supplementary Fig. S2

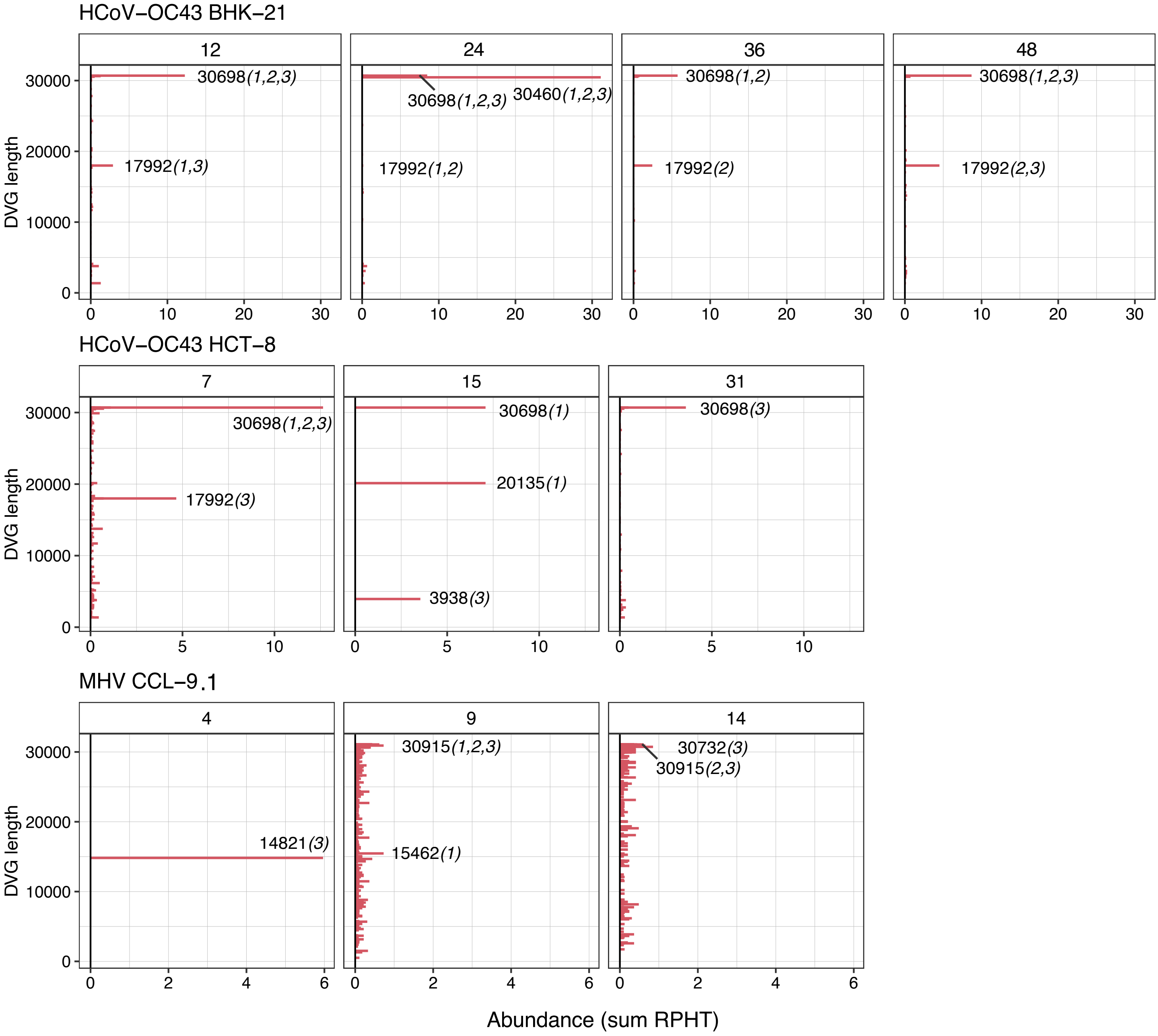
