## Supplementary Table S1 for "Accumulation dynamics of defective genomes during experimental evolution of two betacoronaviruses"

**Table S1.** Statistics describing the variation in MOI for each experimental evolution lineage. Median values ±IQR are shown. Last column shows the fold difference between high and low MOIs (averaging lineages).

| Virus | Cells | Lineage | Low MOI | High MOI | Log$_2$-fold |
|---|---|---|---|---|---|
| HCoV-OC43 | BHK-21 | 1 | $3.74\times10^{-4} \pm2.50\times10^{-2}$ | 21.25 ±82.13 | 16.0 |
| | | 2 | $7.13\times10^{-4} \pm2.50\times10^{-2}$ | 31.25 ±208.50 | |
| | | 3 | $1.50\times10^{-4} \pm1.56\times10^{-2}$ | 25.00 ±145.75 | |
| | HTC-8 | 1 | $5.81\times10^{-5} \pm2.50\times10^{-3}$ | 0.88 ±4.76 | 12.6 |
| | | 2 | $1.21\times10^{-4} \pm3.37\times10^{-3}$ | 0.63 ±2.22 | |
| | | 3 | $3.00\times10^{-3} \pm2.96\times10^{-3}$ | 0.75 ±5.07 | |
| MHV | CCL-9.1 | 1 | $2.65\times10^{-6} \pm8.26\times10^{-6}$ | $6.96\times10^{-3}$ ±0.28 | 12.0 |
| | | 2 | $2.59\times10^{-6} \pm1.79\times10^{-5}$ | $1.06\times10^{-2} \pm9.02\times10^{-2}$ | |
| | | 3 | $8.69\times10^{-6} \pm1.10\times10^{-4}$ | $1.09\times10^{-2} \pm6.62\times10^{-2}$ | |
