## Supplementary Table S2 for "Accumulation dynamics of defective genomes during experimental evolution of two betacoronaviruses"

***Table S2.*** Fitting of viral load data to the best-fitting ARIMA(*p, d, q*)[1] model based on the minimum BIC criterium. Errors represent ±1 SE.

| **Virus** | **Cells** | **MOI** | **Lineage** | **ARIMA** | **BIC** | ***Intercept*** | **Slope** | **Moving average** | **Autocorrelation 1** | **Autocorrelation 2** |
|---|---|---|---|---|---|---|---|---|---|---|
| HCoV-OC43 | BHK-21 | High | 1 | 0, 0, 1 | 64.54 | 8.717 ±0.143 | 0.015 ±0.005 | 0.319 ±0.118 | | |
| | | | 2 | 1, 0, 0 | 142.25 | 11.779 ±0.386 | 0.025 ±0.013 | | 0.394 ±0.138 | |
| | | | 3 | 0, 0, 0 | 232.22 | 18.415 ±0.612 | 0.041 ±0.022 | | | |
| | | Low | 1 | 1, 0, 0 | 546.11 | | −0.820 ±1.228 | | 0.821 ±0.076 | |
| | | | 2 | 2, 0, 1 | 143.65 | 13.336 ±0.465 | 0.017 ±0.016 | 0.565 ±0.161 | −0.302 ±0.176 | 0.565 ±0.136 |
| | | | 3 | 2, 0, 0 | 243.91 | 18.850 ±1.352 | 0.043 ±41.346 | | 0.268 ±0.268 | 0.284 ±0.146 |
| | HCT-8 | High | 1 | 0, 0, 0 | 631.98 | | −48.459 ±41.346 | | | |
| | | | 2 | 0, 0, 0 | 104.77 | 14.737 ±0.365 | −0.097 ±0.020 | | | |
| | | | 3 | 0, 0, 0 | −22.13 | 6.895 ±0.050 | −0.018 ±0.003 | | | |
| | | Low | 1 | 0, 0, 1 | 640.54 | | −635.858 ±74.643 | 0.720 ±0.137 | | |
| | | | 2 | 1, 0, 0 | 346.36 | | −6.764 ±0.807 | | 0.462 ±0.462 | |
| | | | 3 | 2, 0, 0 | 374.66 | | −4.469 ±2.255 | | 0.425 ±0.425 | 0.335 ±0.183 |
| MHV | CCL-9.1 | High | 1 | 0, 0, 0 | 66.58 | 7.8304 ±0.367 | −0.036 ±0.030 | | | |
| | | | 2 | 0, 0, 0 | 33.85 | 5.584 ±0.174 | 0.007 ±0.014 | | | |
| | | | 3 | 0, 0, 0 | −15.11 | 3.793 ±0.002 | 0.057 ±0.005 | | | |
| | | Low | 1 | 1, 0, 0 | 446.34 | | −789.109 ±330.142 | 0.781 ±0.130 | | |
| | | | 2 | 1, 0, 0 | 446.46 | | −697.742 ±338.711 | 0.774 ±0.142 | | |
| | | | 3 | 1, 0, 0 | 446.15 | | −866.648 ±317.089 | 0.779 ±0.125 | | |

[1]ARIMA parameters: $p \geq 0$: order of the autoregressive model (number of time lags); $d \geq 0$: is the differencing order; $q \geq 0$: order of the moving-average model.
